# Redistribution of excitation energy between two photosystems during light-shade adaptation in marine diatoms: State conversion of light-harvesting complexes

**DOI:** 10.1101/2025.11.13.688172

**Authors:** Natsuko Inoue-Kashino, Minoru Kumazawa, Shimpei Aikawa, Kumiko Fujimoto-Omori, Tomoko Ishihara-Masunaga, Sakae Kudoh, Kazuhiko Satoh, Yuichiro Takahashi, Kentaro Ifuku, Yasuhiro Kashino

**Affiliations:** Graduate School of Science, University of Hyogo, 3-2-1 Koto, Kamigohri, Ako-gun, Hyogo 678-1297, Japan; Graduate School of Agriculture, Kyoto University, Kyoto 606-8502, Japan; Biological Resources and Post-Harvest Division, Japan International Research Center for Agricultural Sciences (JIRCAS), 1-1 Ohwashi, Tsukuba, Ibaraki 305-8686, Japan; National Institute of Polar Research, 10-3 Midori-cho, Tachikawa, Tokyo 190-8518, Japan; Department of Polar Science, The Graduate University for Advanced Studies, 10-3 Midori-cho, Tachikawa, Tokyo 190-8518, Japan; Research Institute for Interdisciplinary Science, Okayama University, 3-1-1 Tsushima-Naka, Kita-ku, Okayama 700-8530, Japan

**Keywords:** Light acclimation, Marine diatom, Photosynthesis, Photosystem, State transitions

## Abstract

Marine diatoms effectively photosynthesize by acclimating to the wide range of growth irradiance in the ocean. Using a pennate diatom *Phaeodactylum tricornutum* and a centric diatom *Chaetoceros gracilis*, we evaluated differences in the photosynthetic machinery under a wide range of growth irradiances. The chlorophyll *a*-specific amounts of the major accessory pigments remained relatively constant irrespective of growth irradiance in both diatoms. However, fluorescence spectra at 77K differed drastically depending on the growth irradiance: In *P. tricornutum*, fluorescence from photosystem II was dominant in high-light-grown cells and negligible in low-light-grown cells, while in *C. gracilis*, the opposite trend was observed. These drastic changes in fluorescence spectra were slow processes. The amounts of the two reaction centers, as assessed by specific antibodies and absorption changes in P700, remained almost constant under different irradiances. These results indicate that under dim growth irradiance, more excitation energy is diverted to photosystem I in the pennate diatom, and to photosystem II in the centric diatom. Therefore, the light-harvesting antennas balance excitation energy distribution by changing their association between photosystems I and II in different manners between *P. tricornutum* and *C. gracilis*, depending on irradiance. This phenomenon is similar to state transitions, but differs in its magnitude and duration. Differences in the preference of energy distribution in the two diatoms suggest that the dynamic ‘state conversion’—an antenna rearrangement during the long-term acclimation process in diatoms—is the species-specific strategy to achieve effective photosynthesis under the wide range of growth irradiances in the ocean.

## INTRODUCTION

Diatoms greatly contribute to the global carbon cycle and contribute to regulation of the global climate because photosynthesis (carbon assimilation) in diatoms accounts for approximately 20% of global annual primary production (Nelson et al., 1995). There are more than 100,000 species of diatoms, making them the most abundant type of microalgae in terms of species number (Sims et al., 2006). Because of their high photosynthetic productivity and their ability to produce valuable metabolites, there is increasing industrial interest in diatoms for production of biodiesel and other valuable biological materials (Ramachandra et al., 2009; Tokushima et al., 2016; Vinayak et al., 2015; Zaslavskaia et al., 2001).

In the ocean, diatoms are distributed from the sea surface to the bottom of the euphotic zone (e. g., (Hashihama et al., 2010)). The irradiance in the diatom’s habitat depends on the depth. Because diatoms move up and down in the water column, they should be able to acclimate so as to regulate photosynthetic performance under different irradiances. It was reported that, in the marine diatom *Skeletonema costatum*, the antenna size based on the reaction center of PSI (P700) decreased from 1,100 to 650 molecules of chlorophyll (Chl) *a* per P700 and the number of cellular P700 increased when growth light increased from 0.7 to 130 µmol photons·m^−2^·s^−1^ (Falkowski and Owens, 1980). The opposite change was observed in the marine chlorophyte *Dunaliella tertiolecta*, in which the antenna size increased from 360 to 530 Chl *a* molecules per P700 and the number of P700 decreased from 61 x 10^5^ to 15.8 x 10^5^ P700 per cell when growth irradiance increased from 2 to 400 µmol photons·m^−2^·s^−1^ (Falkowski and Owens, 1980). Because the reaction rate of PSI is approximately 50 times faster than that of PSII (Falkowski et al., 1981), the regulation of PSI–PSII stoichiometry is optimized to balance photosynthetic electron transport when ambient irradiance changes. Although it was shown that the PSII/PSI ratio does not change markedly in response to changes in irradiance in *D. tertiolecta* (Falkowski and Owens, 1980), changes in the PSII/PSI ratio in response to changes in irradiance can be large in diatoms, and can show opposite patterns depending on the species (Hihara and Sonoike, 2001; Strzepek and Harrison, 2004). In addition to the regulation of photosystems ratio, dissipation of excess light energy as heat using Lhcx proteins and some types of light-harvesting proteins (Beer et al., 2006; Zhu and Green, 2010) contributes to the photoprotection under fluctuating irradiance and high irradiance in diatoms ((Bailleul et al., 2010; Zhang et al., 2024), see (Goss and Lepetit, 2015) for review). Dissipating mechanism is still under debate but is known to be coupled with xanthophyll cycle called as diadinoxanthin-diatoxanthin cycle in diatoms (Kashino and Kudoh, 2003; Olaizola and Yamamoto, 1994) (see (Goss and Lepetit, 2015) for review).

Precise physiological analyses are required to understand the productive photosynthesis of diatoms and the mechanism by which photosynthesis is regulated under the wide range of ambient growth irradiances in the ocean, from the surface to the deeper layers. Towards the general understanding of diatoms and the development of genetic manipulation techniques, the entire genome sequence has been determined for over ten diatom species (Falciatore et al., 2020). Despite progress in genomic analysis and the growing interest in understanding and exploiting diatoms, the hard amorphous silica shell of the diatom cell had been proved to be a barrier for biochemical analyses. To overcome this problem, we developed a simple freeze–thaw method to isolate intact thylakoids (Ikeda et al., 2005, 2008). This method has allowed us to conduct in-depth analyses of the biochemical properties of photosystems I and II (PSI and PSII) in diatoms (Ikeda et al., 2008; Nagao et al., 2010) and resulted in precise understandings of the structures of both photosystems (Nagao et al., 2019, 2020, 2022).

Accordingly, it was revealed that, in a marine centric diatom, *Chaetoceros gracilis*, the monomeric form of PSI harbors 24 antenna subunits of light-harvesting chlorophyll-binding protein complexes (LHCs) (Macpherson and Hiller, 2003), which are also named as fucoxanthin chlorophyll-binding protein complexes (FCP) (Xu et al., 2020). The supercomplex contains 326 Chl *a*, 34 Chl *c*, and 155 carotenoids. Nagao et al. reported a smaller PSI-FCPI supercomplex from the same *C. gracilis* (Nagao et al., 2020); PSI binds 16 different FCPs and contains 222 Chl *a*, 54 Chl *c*, and 105 carotenoids, which suggests peripheral FCPs were detached from the large PSI-FCPI supercomplex. PSII-FCPII supercomplex is a homodimer, and each monomer binds 11 FCPs and 144 or 145 Chls, 73 or 72 carotenoids (Nagao et al., 2022; Pi et al., 2019). It is noteworthy that both PSI and PSII carry large antenna pigments. Genomic analyses have revealed that diatoms have more than 40 genes encoding FCPs, which are categorized into six subfamilies: Lhcr, Lhcf, Lhcx, Lhcz, Lhcq, and CgLhcr9 homologs (Kumazawa et al., 2022). The Lhcr subfamily is found in a wide variety of red algal lineages. *C. gracilis* genome contains 9 Lhcr subfamily genes (Lhcr1–3, Lhcr5–8, Lhcr10, and Lhcr17), and PSI in *C. gracilis* associates 8 Lhcr except for Lhcr17 assigned in PSII-FCPII (Kumazawa et al., 2022). The Lhcz subfamily is an independent clade of the Lhcr subfamily. The Lhcf subfamily includes FCPs associated with PSII (Nagao et al., 2020). The Lhcx subfamily participates in energy-dependent NPQ (qE) to dissipate excess light energy as heat. The Lhcq subfamily comprises the peripheral belt of FCPs in the PSI-FCPI supercomplex of *C. gracilis* (Nagao et al., 2020; Xu et al., 2020). CgLhcr9 forms an independent clade in phylogenetic tree, and is found in the PSI-FCPI of *C. gracilis*.

In the present study, to reveal light-acclimation mechanisms in diatoms, we analyzed changes in the composition of photosynthetic systems and photosynthetic pigments in two diatom species under a wide range of growth irradiances (3–320 μmol photons m^−2^ s^−1^). The two marine diatoms were the pennate diatom *Phaeodactylum tricornutum* and the centric diatom *Chaetoceros gracilis* for which highly efficient transformation systems had been established by our group (Ifuku et al., 2015; Miyahara et al., 2013).

## RESULTS

### Changes in Amounts of Photosynthetic Pigments Under a Range of Irradiances

Fucoxanthin and Chl *c*, the major light-harvesting pigments in diatoms, bind to the FCP (Macpherson and Hiller, 2003). We measured changes in the relative amounts of fucoxanthin and Chl *c* to Chl *a* in *P. tricornutum* and *C. gracilis* cells under different growth light intensities by HPLC analysis. In both species, the molar ratios of fucoxanthin and Chl *c* to Chl *a* did not change in response to different growth irradiances (**Fig. 1**). In *P. tricornutum*, the combined amount (DD+DT) of diadinoxanthin (DD) and diatoxanthin (DT), which are responsible for the photoprotective xanthophyll cycle (Kashino and Kudoh, 2003; Olaizola and Yamamoto, 1994), and the amount of ß-carotene, which also has a role in photoprotection (Scheer, 2003), significantly increased under high light (320 µmol photons·m^−2^·s^−1^). In *C. gracilis*, the increases in the amounts of these photoprotective pigments under elevated irradiances were limited compared with those in *P. tricornutum*. The molar ratio of pheophytin *a*, the primary electron acceptor in PSII, to Chl *a* remained almost constant irrespective of the growth irradiance. The results shown in **Fig. 1** are consistent with previous findings in our laboratory (Ban et al., 2006); in both species, the molar ratios of fucoxanthin and Chl *c* to Chl *a* did not markedly change in response to different growth irradiances, and xanthophyll cycle-related pigments increased under higher growth irradiances. Similar results were reported for *P. tricornutum* (Gundermann et al., 2013; Nagao et al., 2019) although the amount of ß-carotene did not change in (Nagao et al., 2019b).

**Fig. 1.**
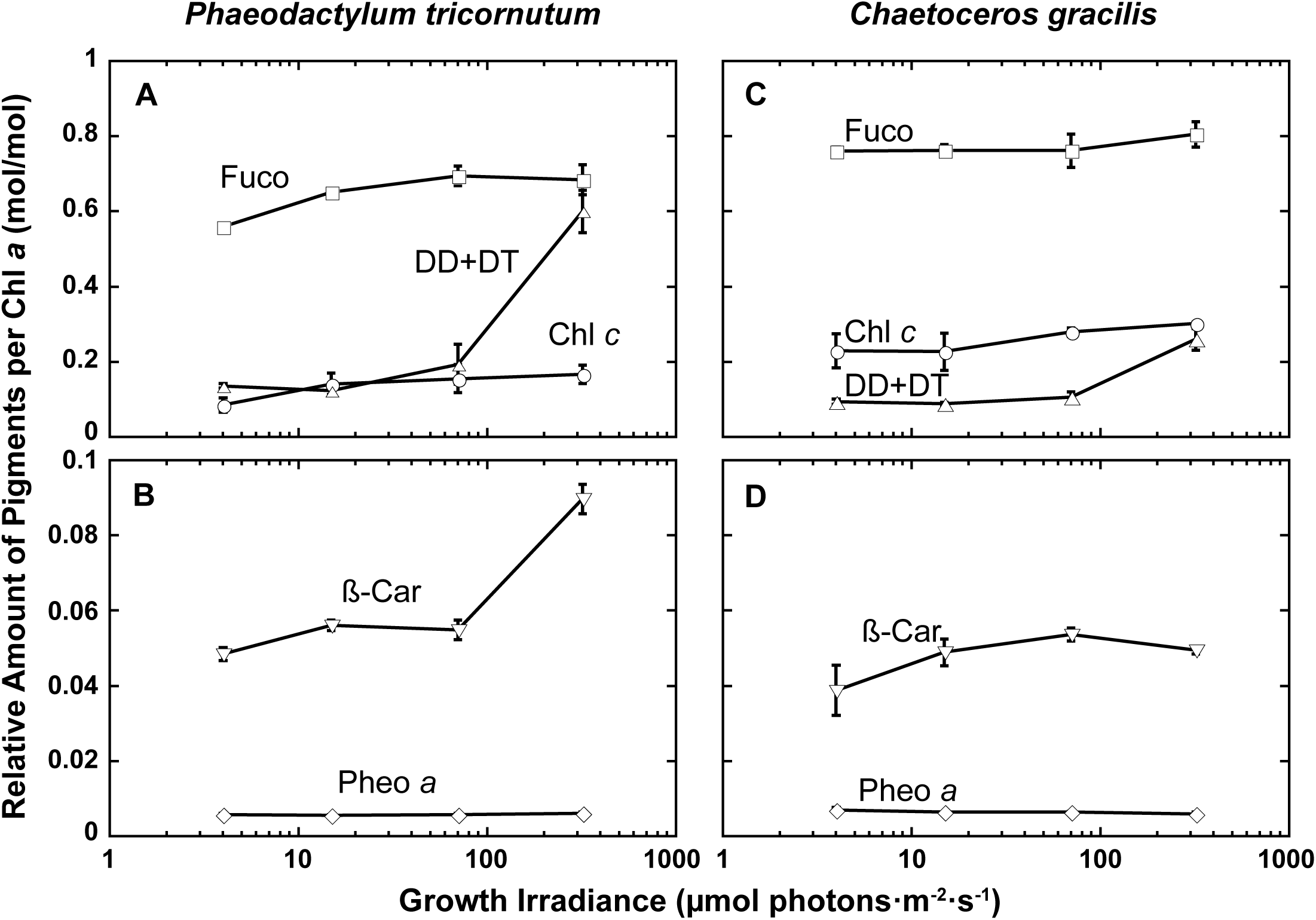
Changes in amount of accessory pigments to Chl *a* in *P. tricornutum* (A and B) and *C. gracilis* (C and D) grown under different growth irradiances. Molar ratios of fucoxanthin (Fuco; squares), chlorophyll *c* (Chl *c*; circles), diadinoxanthin + diatoxanthin (DD+DT; triangles), ß-carotene (ß-Car; inverted triangles), and pheophytin *a* (Pheo *a*; diamonds) to Chl *a* are shown. Error bars (se, *n*=3–4, except lowest irradiance for *P. tricornutum* [*n*=2]) are sometimes smaller than the symbols.

### Purification and Characterization of Photosystem I and II Complexes from *Phaeodactylum tricornutum*

Since the fluorescence observed for PSI and PSII in *C. gracilis* is already reported (Ikeda et al., 2008; Nagao et al., 2007), to assess the fluorescence emitted from PSI and PSII in *P. tricornutum* at 77K, PSI and PSII complexes were purified from *P. tricornutum*. For this purpose, isolated thylakoid membranes were solubilized in 0.5%–3.0% *n*-dodecyl-β-D-maltoside (DDM), 0.5%–2.0% *n*-heptiyl-ß-D-thioglucoside (HTG), and 0.5%–2.0% Triton X-100. The resulting extracts were subjected to sucrose density gradient ultracentrifugation. As shown in **Fig. 2A**, extracts prepared using 3.0% DDM showed superior separation of one brown band and two green Chl-containing bands. The polypeptide profiles of the two green bands F1 and F2 showed typical polypeptide patterns of PSI and PSII, respectively (**Fig. 2B**). Western blot analysis confirmed that fractions F1 and F2 are PSI and PSII complexes, respectively.

**Fig. 2.**
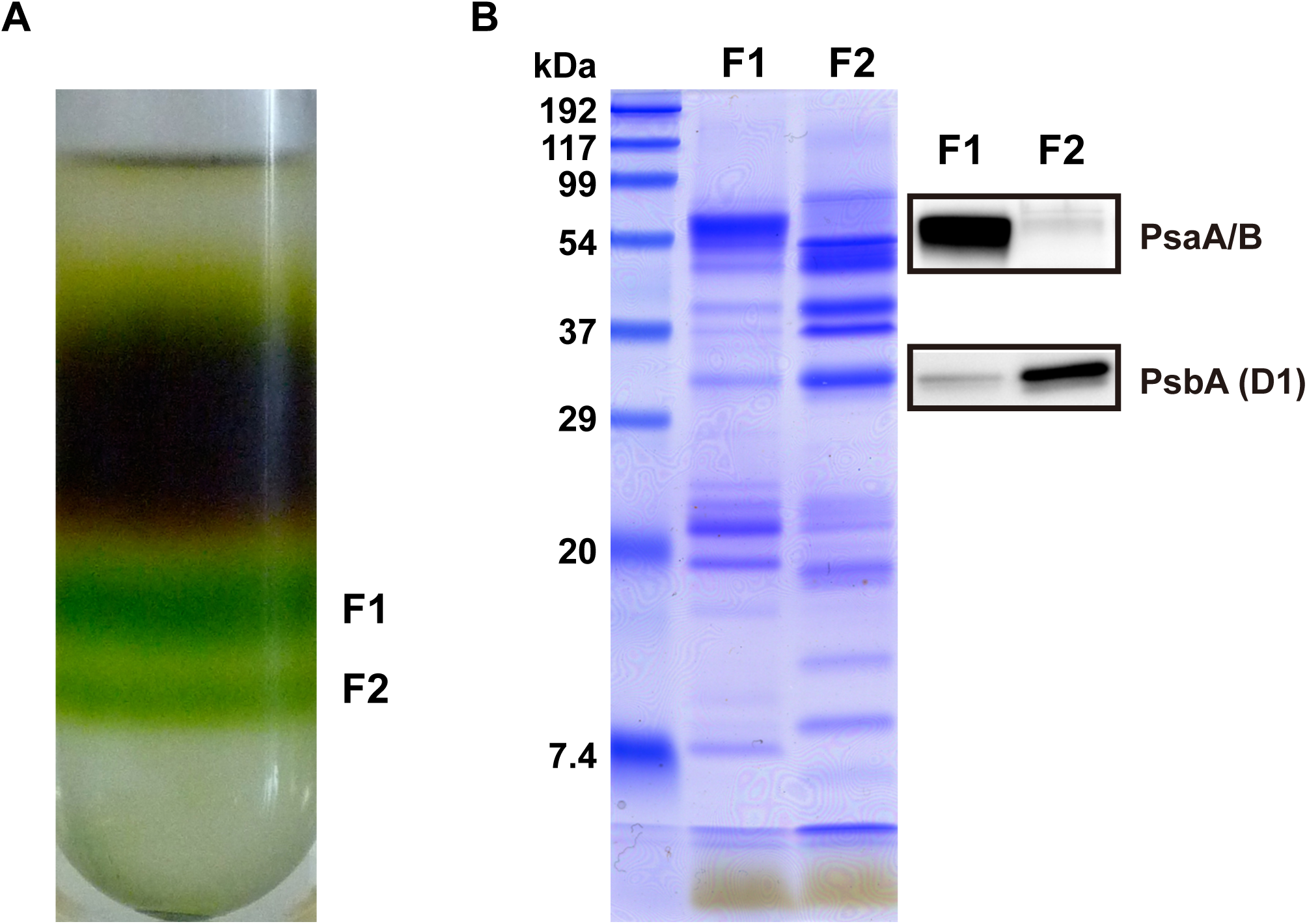
Separation of chlorophyll-binding protein complexes isolated from the pennate diatom *Phaeodactylum tricornutum* by sucrose density gradient centrifugation (A) and polypeptide profiles of resulting F1 and F2 fractions (B). Thylakoid membranes (1.0 mg Chl *a*/mL) were solubilized using 3.0% DDM and subjected to sucrose density gradient centrifugation. Polypeptides in F1 and F2 fractions were visualized by Coomassie staining after SDS-PAGE. Specific antibodies detected PS I reaction center proteins PsaA/B and PS II reaction center protein PsbA. Proteins corresponding to 2 and 0.4 µg Chl *a* were separated by electrophoresis for Coomassie staining and immunostaining, respectively.

Fluorescence spectra of fractions F1 and F2 were recorded at 77K. Fluorescence from PSII (**Fig. 3**, green line) showed a peak at ∼695 nm and a shoulder at ∼685 nm, which are typical of the PSII complex (van Dorssen et al., 1987; Murakami, 1997; Murata, 1969; Murata and Satoh, 1986). A small fluorescence hill at ∼750 nm originates from the satellite vibrational bands of 685 and 695 nm fluorescence (Murata and Satoh, 1986; Rijgersberg and Amesz, 1980). The low ratio of signal to noise of the PSI fluorescence spectrum is ascribed to the low fluorescence yield (**Fig. 3**, red line). However, the peak profile was clearly determined; one peak at ∼713 nm corresponding to fluorescence originating from the PSI complex (Itoh et al., 2004; Murata, 1969; Murata and Satoh, 1986) and small shoulders at ∼685 and ∼695 nm corresponding to the fluorescence from a trace amount of contaminated PSII with higher fluorescence yield (van Dorssen et al., 1987; Murakami, 1997; Murata, 1969; Murata and Satoh, 1986).

**Fig. 3.**
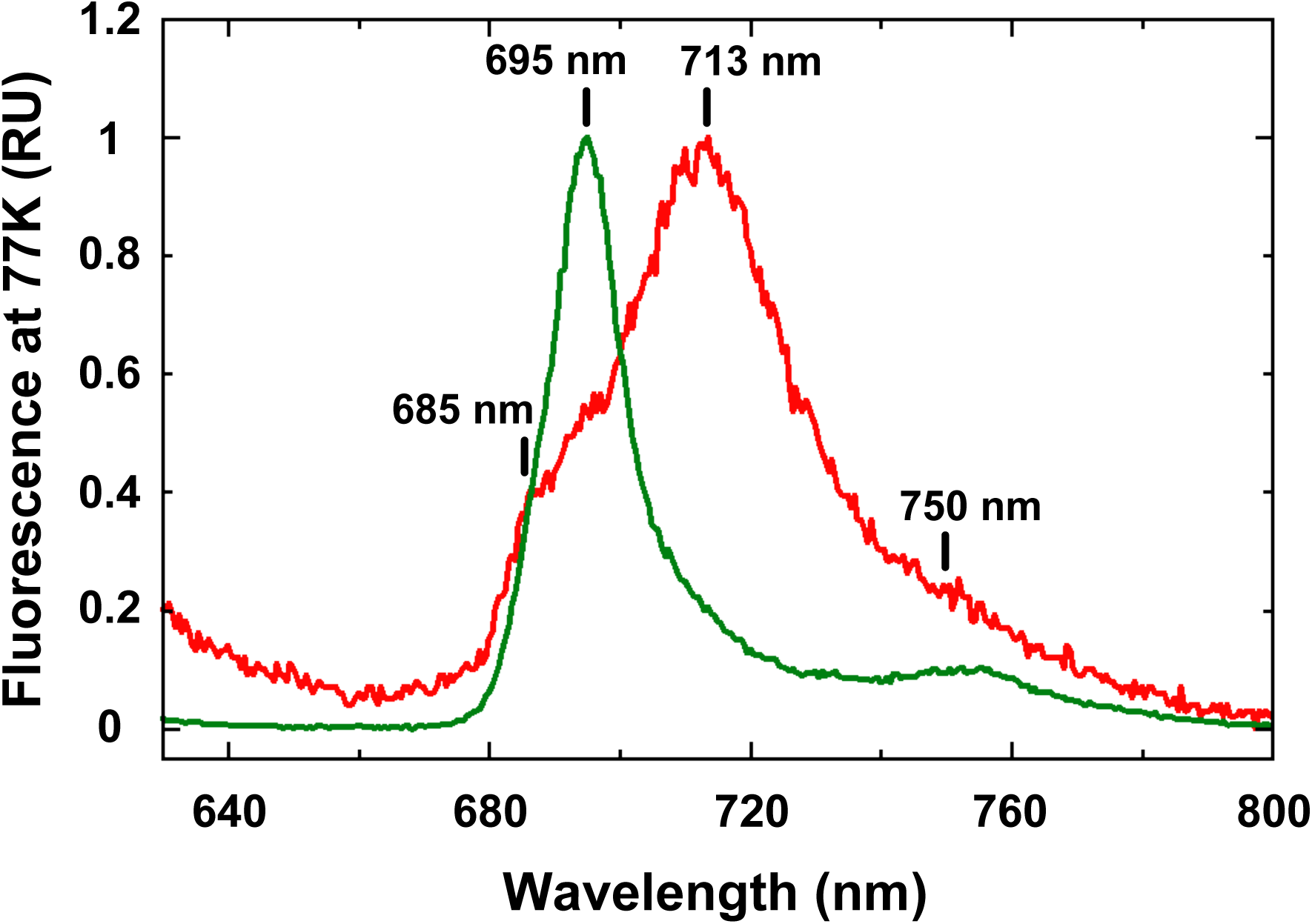
Fluorescence spectra at 77K for photosystems I (red) and II (green) purified from *Phaeodactylum tricornutum*. Photosystems I and II correspond to F1 and F2 in Fig. 2, respectively. For clarity, spectra were normalized at highest peaks.

### Fluorescence Spectra Emitted from Cells Grown Under a Wide Range of Irradiances

We examined the effect of growth irradiance from low to high (3, 15, 70, 320 µmol photons·m^−2^·s^−1^) on the 77K fluorescence spectrum. *P. tricornutum* cells grown under moderate and high irradiances showed a peak profile typical of diatoms and brown algae (Sugahara et al., 1971), with two fluorescence peaks at ∼690 nm and 705–717 nm (**Fig. 4A**). The fluorescence at ∼690 nm (FL690 hereafter) can be emitted from PSII (**Fig. 3**) (Sugahara et al., 1971). When grown under dim growth irradiance (3 µmol photons·m^−2^·s^−1^), most fluorescence was emitted at longer wavelength (715-717 nm; FL715 hereafter) that can be assigned to PSI fluorescence, while FL690 was negligible (**Fig. 4A**, blue line). Under higher growth irradiance, FL690 increased significantly and became larger than FL715 at the highest irradiance tested (320 µmol photons·m^−2^·s^−1^) (**Fig. 4A**, red line).

**Fig. 4.**
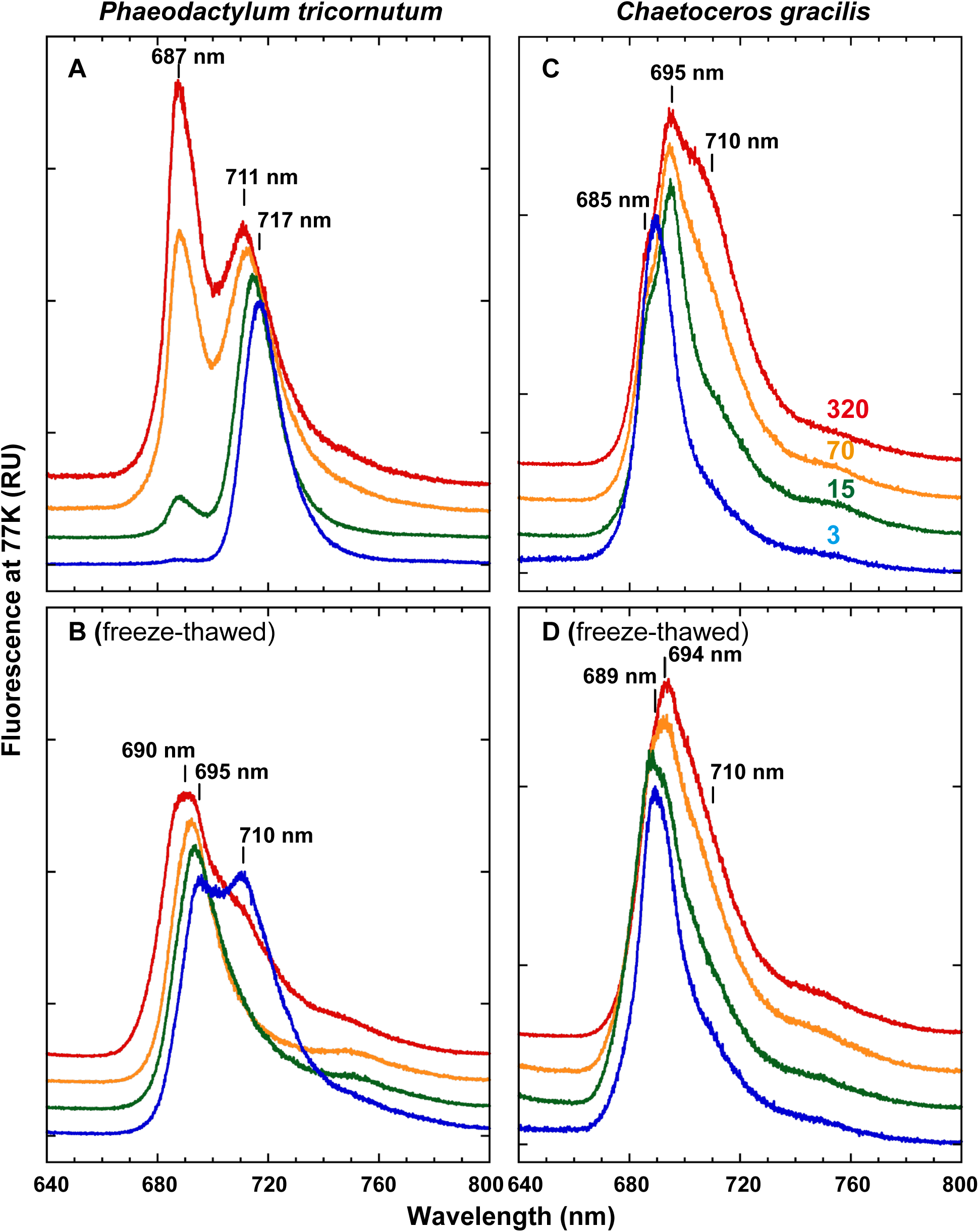
Fluorescence spectra at 77K obtained for *P. tricornutum* (A and B) and *C. gracilis* (C and D). Spectra were measured immediately after cells were harvested (A and C) or after cells were freeze–thawed once (B and D). Cells were grown under light conditions of 3 (blue line), 15 (green line), 70 (orange line) or 320 (red line) µmol photons·m^−2^·s^−1^. Spectra were normalized at 710 nm (A) and 690 nm (B), or at the highest peaks (C and D). Typical traces are drawn from three to four independent assays.

The fluorescence spectrum of *C. gracilis* at 77K did not show distinct separated peaks from PSI and PSII, whereas there was one peak mainly from PSII with a shoulder from PSI (**Fig. 4C**). This is because PSI emits fluorescence at shorter wavelengths, peaking at 710 nm (FL715) (Ikeda et al., 2008), while the 695 nm fluorescence (FL690) from PSII is similar to that in other organisms (Nagao et al., 2010). FL690 appeared dominant under low-irradiance conditions, while FL715 appeared dominant under high-irradiance conditions (**Fig. 4C**).

To elucidate the changes in the ratio of FL690 and FL715 quantitatively, the fluorescence spectra were deconvolved into two fluorescence components (**Fig. 5A–D**), and the proportions of FL690 and FL715 were estimated (**Fig. 6**). Both strains showed significant changes in FL690 to FL715 ratio when exposed to different growth irradiance conditions but exhibited opposite trends (**Fig. 6**). Such large changes in the ratio of PSII to PSI have not been reported previously, even for diatoms (Falkowski and Owens, 1980; Falkowski et al., 1981; Smith and Melis, 1988; Strzepek and Harrison, 2004). Although the 77K fluorescence spectrum is frequently used to assess the relative amounts of PSI and PSII (Mimuro, 2004; Murakami, 1997; Murata and Satoh, 1986), it is also used to evaluate the ratio of light-energy distribution between the two photosystems (Lamb et al., 2018). Large changes in the proportion of PSI to Chl have not been reported until now. As described above, the antenna complexes in diatoms are huge. Thus, the re-localization of antennas in response to changes in growth irradiance could be observed as changes in the PSI- to PSII-fluorescence ratio, because the distribution of captured light energy in the cell depends on the size of antennas associated with PSI and PSII. In other words, the huge antennas could perturb the proportion of fluorescence components by transferring light energy preferentially to one of the two photosystems. Therefore, the cells were frozen and thawed before measuring fluorescence (**Fig. 4B, D**) to interfere with energy transfer from antennas to photosystems by breaking functional structure of PSI-FCPI and PSII-FCPII supercomplexes. After freeze-thawing, the spectra changed drastically. In *P. tricornutum*, the fluorescence at longer wavelengths (FL715) decreased significantly under all growth-irradiance conditions except the highest irradiance, and fluorescence at shorter wavelengths (FL690) increased under all growth-irradiance except the lowest irradiance (**Fig. 4B**). FL715 was prominent in cells grown under the lowest and highest light intensities (blue and red lines in **Fig. 4B**, respectively). The fluorescence peak of FL690 became broader (**Fig. 4B**) than the corresponding peak measured without freeze–thawing (**Fig. 4A**). This is a result of the contribution of the fluorescence emitted by FCPs, which were detached from PSII-FCPII supercomplex, because the FCPs emit fluorescence at slightly shorter wavelengths (680–685 nm) than that emitted by PSII (695 nm) at 77K (Ikeda et al., 2008; Ishihara et al., 2015; Lepetit et al., 2007). Drastic changes in the 77K fluorescence spectra after freeze–thawing were also observed for *C. gracilis* (**Fig. 4C, D**). FL715 decreased significantly under all growth irradiances except the lowest irradiance (**Fig. 4D**). These results indicated that the changes in the fluorescence spectra in response to different growth irradiances do not reflect significant changes in the proportions of photosystems, but instead reflect the drastic re-localization of light-harvesting antennas (FCPs), although the possibility of the changes in the occurrence of spill-over could not be excluded. The re-localization of light-harvesting antennas in response to irradiance is similar to that during state transitions. However, the response we observed was much slower than state transitions. The spectrum remained almost the same for more than 6 h after switching the irradiance from 3 to 50 µmol photons·m^−2^·s^−1^ (**Fig. 7**).

**Fig. 5.**
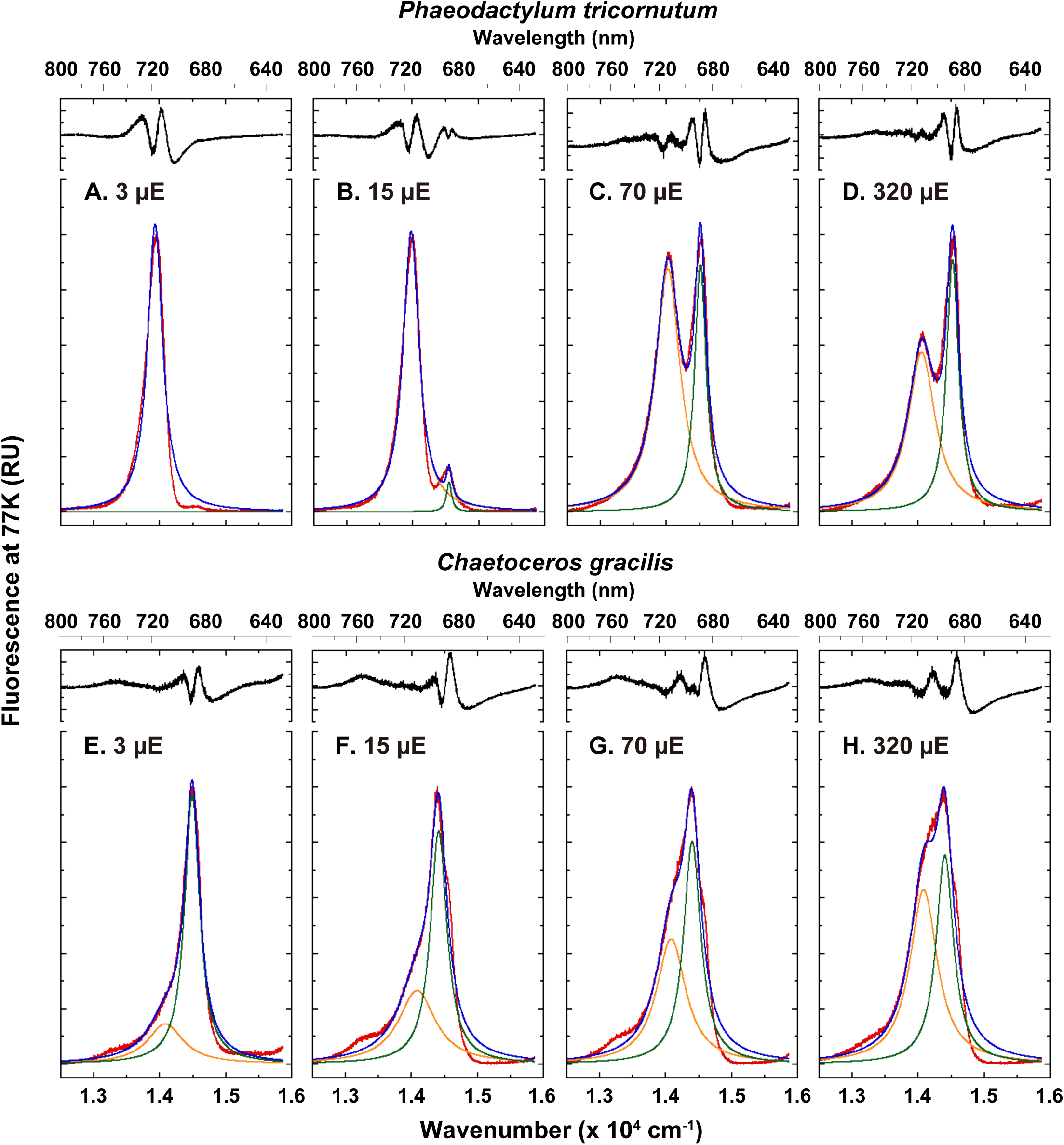
Deconvolution of fluorescence spectra at 77K. Fluorescence spectra for *P. tricornutum* (A–D) and *C. gracilis* (E–H) shown in Fig. 4A and 4C were re-drawn against wavenumber (inverse of wavelength) and fitted to two Lorentzian components. Growth irradiances were 3 (A and E), 15 (B and F), 70 (C and G) or 320 (D and H) µmol photons·m^−2^·s^−1^. Red line, measured spectra; orange line, smaller wavenumber component corresponding to PS I; green line, larger wavenumber component corresponds to PS II; blue line, sum of the two components; black line, delta of measured (red) and fitted (blue) lines.

**Fig. 6.**
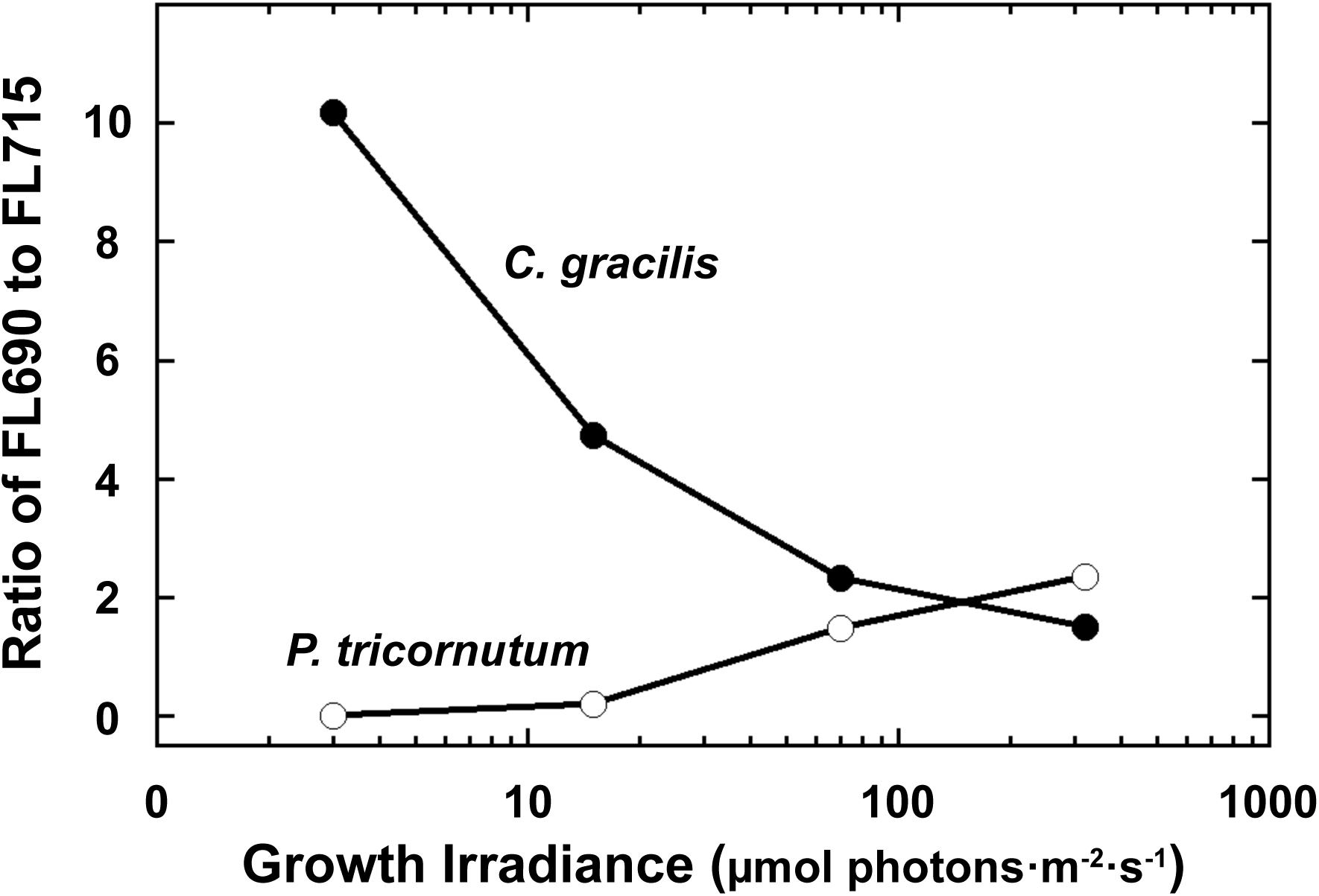
Changes in ratio of two fluorescence components in response to changes in growth irradiance. Relative area of larger wavenumber component (shorter wavelength component: FL690) to smaller component (FL715) in Fig. 5 was plotted against growth irradiance. Open circle; *P. tricornutum*, and closed circle; *C. gracilis*.

**Fig. 7.**
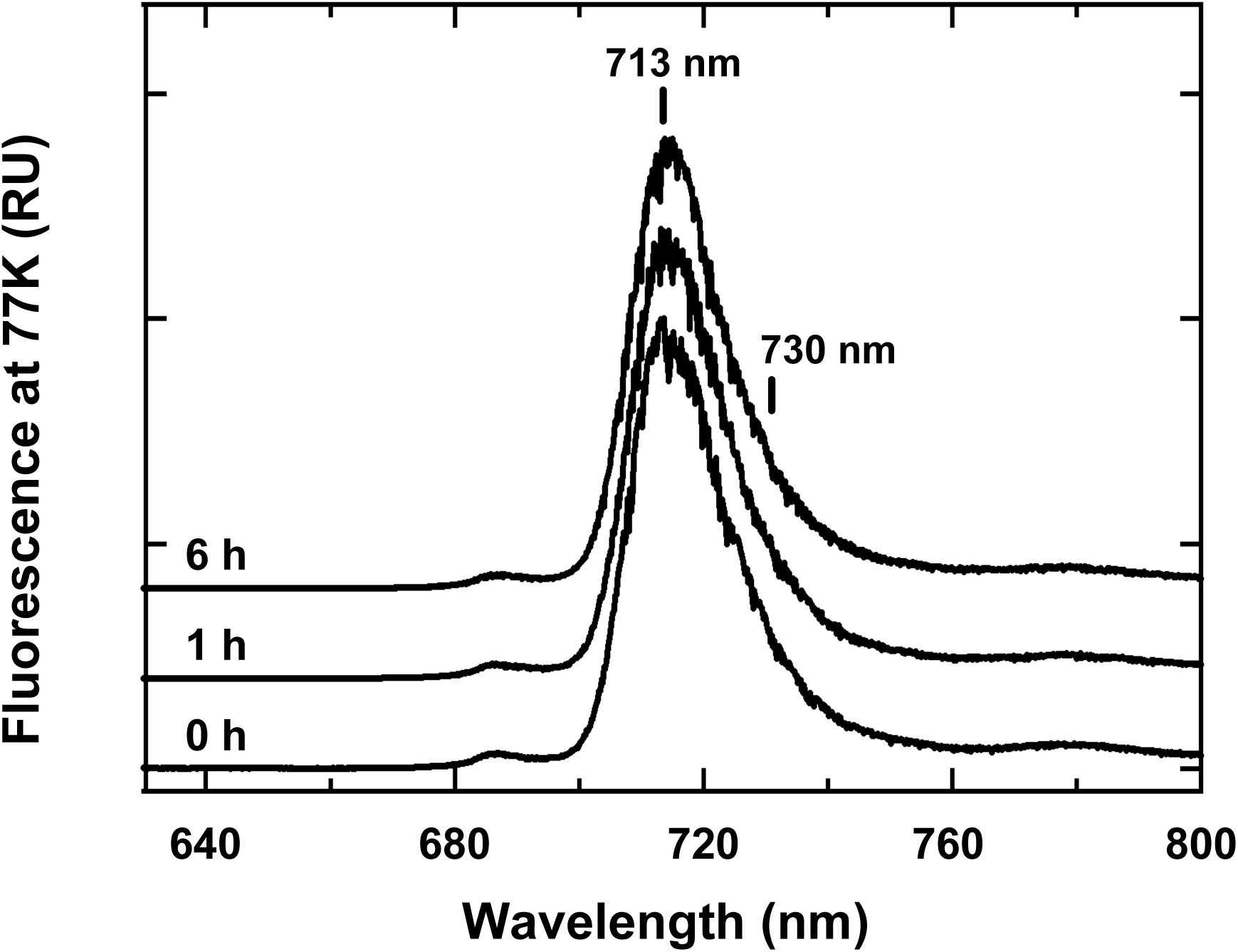
Changes in fluorescence spectra at 77K in *Phaeodactylum tricornutum* during illumination. *P. tricornutum* cells grown under light at 3 µmol photons·m^−2^·s^−1^ were illuminated for 0h, 1 h or 6 h at 200 µmol photons·m^−2^·s^−1^ and their fluorescence spectra were measured at 77K. Chromophores were excited at 430 nm.

### Photosystems Ratios Under Different Growth Irradiances

The amounts of reaction center proteins in two photosystems were assessed using specific antibodies against PsaA/B for PSI and PsbA for PSII (**Fig.** 8). In both diatoms, the relative amounts of PsaA/B and PsbA did not change significantly in response to different growth irradiances.

**Fig. 8.**
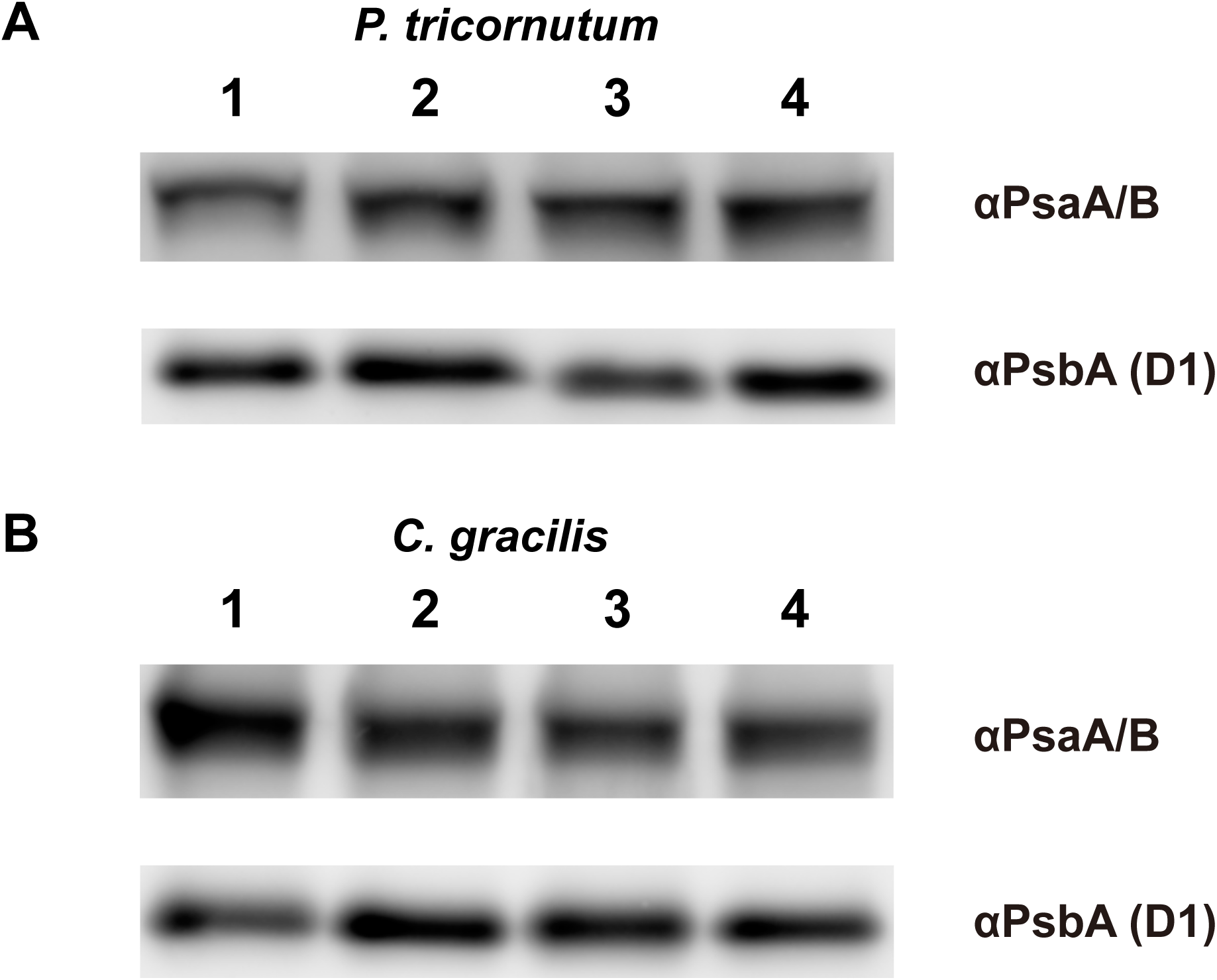
Amount of reaction center proteins in cells grown under different growth irradiances. A, *P. tricornutum*; B, *C. gracilis*. Proteins of cells (0.33 µg Chl *a*) grown under illumination at 3 (lane 1), 15 (lane 2), 70 (lane 3) or 320 (lane 4) µmol photons·m^−2^·s^−1^ were solubilized and subjected to electrophoresis and the reaction center proteins of PS I (PsaA/B) and PS II (PsbA) were detected using specific antibodies.

The amount of photoactive PSI was also assessed by measuring the light-induced oxidation of P700 (**Fig. 9**). P700 was fully photooxidized in the presence of 2,5-dibromo-3-methyl-6-isopropyl-p-benzoquinone (DBMIB), which blocks the linear electron flow from PSII and cyclic electron flow from PSI. Cells illuminated by actinic light for 5 sec fully oxidized P700 and during the subsequent dark period, P700^+^ was slowly re-reduced. Small differences in the amplitude of photoactive P700 were observed between cells grown under high-light and low-light, but these were much smaller than changes in the fluorescence spectra in both diatom cells (**Fig. 9**). The amount of pheophytin *a*, which is a measure of the abundance of PSII, remained almost constant irrespective of the growth irradiance (**Fig. 1**). Thus, it is concluded that the fluorescence spectral changes induced by the cell growth irradiance are not ascribed to changes in the amount of PSI and PSII but to the re-distribution of excitation energy from mobile FCPs between PSI and PSII.

**Fig. 9.**
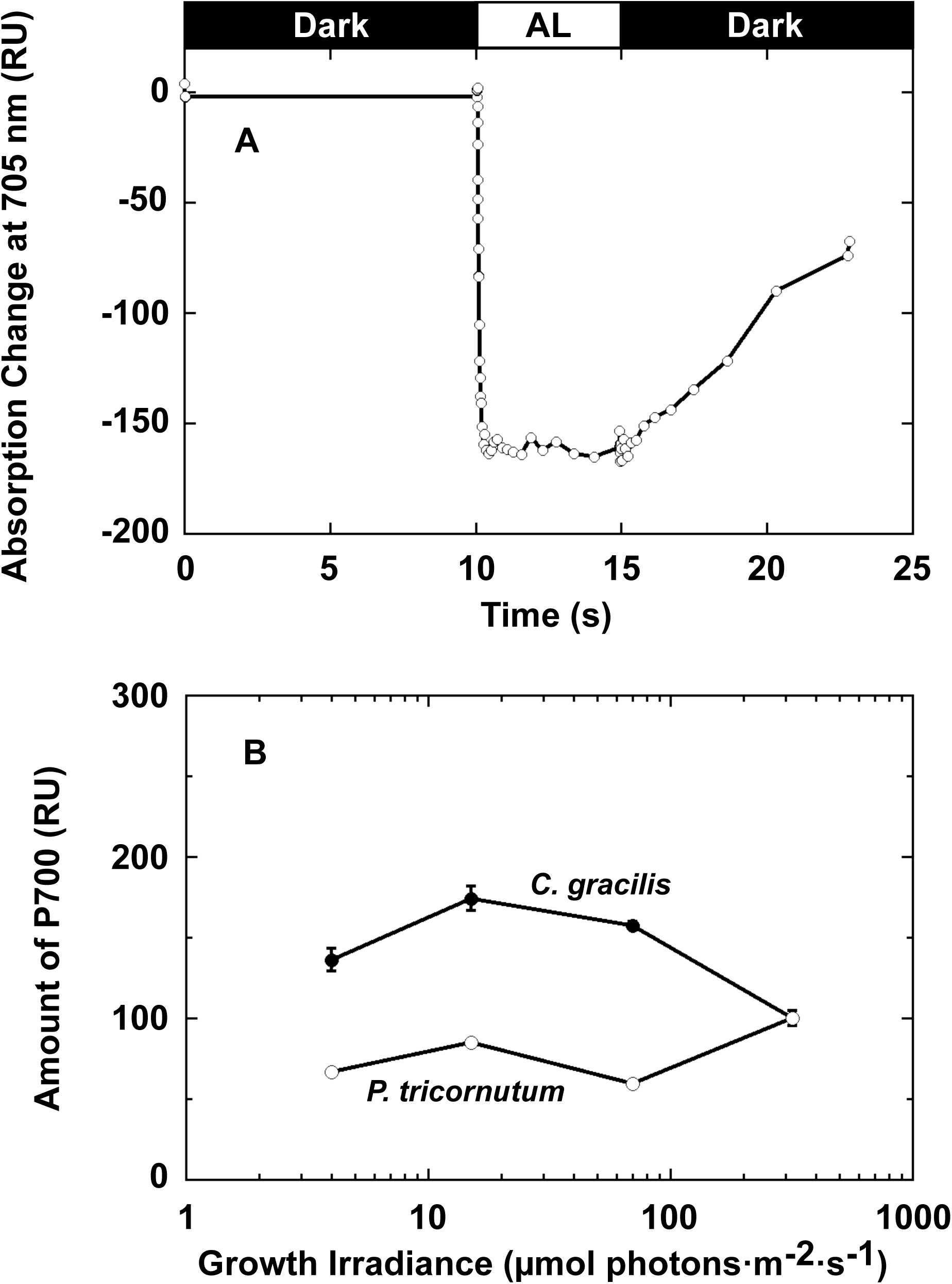
Changes in amount of P700 as measured by changes in absorption. (A) Typical absorption changes of P700 in *C. gracilis* cells monitored at 705 nm with a Joliot-type spectrometer. Cells were grown under light at 70 µmol photons·m^−2^·s^−1^. The reaction mixture contained DBMIB to block linear electron flow from PSII and cyclic electron flow from PSI. (B) Changes in photoactive P700 in *P. tricornutum* (open circle) and *C. gracilis* (closed circle) cells grown under different growth irradiance. Amplitude of photoactive P700 in cells grown under illumination at 320 µmol photons·m^−2^·s^−1^ was set to 100 (*n*=2–9). AL, actinic light.

## DISCUSSION

### Changes in Amounts of Photosynthetic Pigments Under a Wide Range of Irradiances

In land plants, the relative amounts of accessory pigments (such as Chl *b*) increase and Chls are redistributed from reaction center core complexes to light-harvesting protein complexes under low-light conditions, and vice versa under high-light conditions (De la Torre and Burkey, 1990). In light of this well-known pattern of photo-acclimation, it is curious that the molar ratio of fucoxanthin and Chl *c* to Chl *a* did not change significantly in *P. tricornutum* and *C. gracilis* cells grown under different irradiances, as shown in **Fig. 1**. One of the main reasons is that photosynthetic antennas in diatoms are much larger than those in higher plants. For example, in *C. gracilis*, 16 FCPs associated with PSI harbor 128 Chl *a*, 54 Chl *c*, and 85 carotenoid molecules in total in addition to the pigments in the core complex (Nagao et al., 2020), or 24 antenna subunits (FCPs) associated with PSI bind 232 Chl *a*, 34 Chl *c*, and 133 carotenoid molecules in total in addition to the pigments in the core complex (Xu et al., 2020). The FCPs associated with PSII harbor 71 Chl *a*, 35 Chl *c*, and 54 fucoxanthin in addition to the pigments in the core complex (Nagao et al., 2022).

Whereas the ratio of light-harvesting pigments to Chl *a* remained relatively stable in cells grown under different irradiances, the ratio of diadinoxanthin cycle pigments increased as growth irradiance increased, more pronouncedly in *P. tricornutum*. This change is related to the photo-protection mechanisms of the diadinoxanthin cycle through non-photochemical quenching (NPQ) as shown in our previous study (Ban et al., 2006) and other studies (e. g. (Arsalane et al., 1994; Buck et al., 2019; Olaizola et al., 1994)).

### PSI and PSII Complexes in *Phaeodactylum tricornutum*

Despite the long history of *P. tricornutum* as a model diatom and significant efforts by many researchers, the purification of PSI and PSII core complexes from *P. tricornutum* has been unsuccessful, mainly because of the hard silica shell surrounding the cells. We developed a method to obtain intact thylakoid membranes from *C. gracilis* by a simple freeze–thaw process (Ikeda et al., 2005, 2008) and successfully purified PSI (Ikeda et al., 2005, 2008) and PSII (Nagao et al., 2007, 2010) complexes. However, this method was not successful in isolating PSI and PSII complexes from *P. tricornutum*. Instead, we used a method that was developed to prepare thylakoid membrane from cyanobacteria (Kashino et al., 2002) to isolate intact thylakoids from *P. tricornutum* cells. Using these thylakoids, PSI and PSII complexes were purified after solubilization by a mild detergent (DDM) followed by sucrose density gradient ultracentrifugation (**Fig. 2**). To our knowledge, these are the most purified complexes from *P. tricornutum* judging from the polypeptide patterns. Structural analysis of photosystems from *P. tricornutum* is awaited after the reports of structures of photosystems from centric diatoms, *C. gracilis* (Nagao et al., 2022, 2020, 2019; Xu et al., 2020), *Thalassiosira pseudonana* (Feng et al., 2025, 2023), and *Cyclotella meneghiniana* (Zhao et al., 2023).

Using the highly purified PSI and PSII complexes, fluorescence spectra of PSI and PSII at 77 K were obtained (**Fig. 3**). A fluorescence peak at wavelengths longer than ∼705 nm is usually emitted from PSI (Murata et al., 1966; Murata and Satoh, 1986). While this holds true for many photosynthetic organisms, in *P. tricornutum*, both the long and short wavelength fluorescence peaks at 77K have been reported to originate from PSII (Brown, 1967; Fujita and Ohki, 2004; Goedheer, 1973; Shimura and Fujita, 1973). This assignment is based on the observation that, different from many other photosynthetic organisms, *P. tricornutum* emits fluorescence at ∼710 nm even at room temperature, which is induced when *P. tricornutum* is grown under weak red light. In many photosynthetic organisms, PSI does not emit fluorescence at room temperature, but it does emit fluorescence at low temperatures such as 77K (Murata and Satoh, 1986). The wavelength of this room temperature fluorescence is almost the same as that measured at 77K. Furthermore, the fluorescence transient at 707 nm at room temperature is consistent with the oxidation–reduction reaction of Q_A_ in PSII (Shimura and Fujita, 1973). More recent studies supported that this long-wavelength fluorescence (F710) was emitted from FCP (Lhcf15), functionally connected to PSII (Herbstová et al., 2015). However, it was also reported that F710 disappeared quickly *in vitro* (Brown, 1967; Fujita and Ohki, 2004; Herbstová et al., 2015; Sugahara et al., 1971). Also, the fluorescence yield at 77K was shown to be approximately 24 times (Shimura and Fujita, 1973) or 7–10 times (Brown, 1967) higher than that at room temperature. Therefore, F710 could not be observed in PSI and PSII complexes purified *in vitro* in this work (**Fig. 3**).

The wavelength of the fluorescence peak at 77 K for the PSI complex purified in this work was ∼713 nm (**Fig. 3**), consistent with that of PSI complexes in many other photosynthetic organisms (Itoh et al., 2004; Murata, 1969; Murata and Satoh, 1986). The fluorescence profile of purified PSII at 77 K (**Fig. 3**) was the same as those reported from other photosynthetic organisms (van Dorssen et al., 1987; Murata, 1969; Murata and Satoh, 1986). The fluorescence profiles of PSI and PSII in *C. gracilis* were reported (Ikeda et al., 2008; Nagao et al., 2010).

### Proportions of Photosystems I and II in Cells Grown Under a Range of Irradiances

The amounts of PSI and PSII complexes are relatively low compared with the amount of FCPs in diatom cells. Therefore, the fluorescence spectrum at 77K was measured to assess the relative amounts of PSI and PSII (van Dorssen et al., 1987; Murakami, 1997; Murata, 1969; Murata and Satoh, 1986) after confirming the fluorescence profiles of PSI and PSII in *P. tricornutum* (**Fig. 3**). Drastic changes in the proportions of two fluorescence components, FL690 and FL715, were observed in both diatoms (**Figs 3, 4, and 5**). Considering the significant changes in the ratio of FL690 to FL715, the proportions of PSI and PSII appeared to change markedly depending on the growth irradiance. In *P. tricornutum*, the ratio of FL690 to FL715 was ∼2.3 in cells grown under 320 µmol photons·m^−2^·s^−1^, despite the trace amount of FL690 in cells grown under 3 µmol photons·m^−2^·s^−1^. Changes in the ratios of PSII to PSI depending on irradiance have been reported for some other microalgae; e.g., 3.9 (low light: LL) and 1.3 (high light: HL) in the diatom *Cylindrotheca fusiformis* (Smith and Melis, 1988), 2.2 (LL) and 1.1 (HL) in the diatom *Thalassiosira costatum* (Falkowski and Owens, 1980), 8.8 (LL) and 11 (HL) in the oceanic diatom *Thalassiosira oceania* (Strzepek and Harrison, 2004), 1,4 (LL) and 4.2 (HL) in the coastal diatom *Thalassiosira weissflogii* (Strzepek and Harrison, 2004), and 0.91 (LL) and 0.83 (HL) in the green alga *Dunaliella tertiolecta* (Falkowski et al., 1981). However, these changes in the ratios of PSII to PSI in response to different light intensities were not as large as the changes in the ratio of FL690 to FL715 observed in this work. Furthermore, the changes in the amounts of PSI reaction center proteins PsaA/B and PSII reaction center protein PsbA (**Fig. 8**), and the amount of photoactive P700 (**Fig. 9**) were small compared with the changes in fluorescence spectra in **Figs 4–6**. The amount of pheophytin, the primary electron acceptor in PSII, also remained relatively constant under different growth irradiances (**Fig. 1**).

The fluorescence spectra changed markedly after cells were subjected to the freeze–thaw process (**Fig. 4C, D**). In another study, thylakoids from Ochrophyte (Chromophyte) algae (*P. tricornutum* and *Vischeria helvetica*) isolated without using high-osmotic and high-ionic strength medium showed a loss of long-wavelength fluorescence, resulting in a significant change in fluorescence spectra (Chrystal and Larkum, 1988). The high-osmotic and high-ionic strength medium that retained long-wavelength fluorescence was almost identical to that used to stabilize the functional linkage between phycobilisome and PSII core in cyanobacteria (Dilworth and Gantt, 1981; Katoh and Gantt, 1979; Kura-Hotta et al., 1986). These results illustrated that the changes in fluorescence spectra by the freeze–thaw process (**Fig. 4C, D**) can be caused by dissociation of functional coupling between core complexes and the peripheral antenna system. This means that once the antenna system functionally dissociated from the reaction center as a result of freeze–thawing, it emitted fluorescence rather than transferring excitation energy to the reaction center.

Considering the results described here and the huge antenna size in diatoms, it could be assumed that changes in the relative fluorescence amplitude of FL690 and FL715 were the consequences of flexible re-localization of antennas between PSI and PSII in response to changes in growth irradiance. This is because the distribution of captured light energy in the cell depends on the size of antennas associated with PSI and PSII.

Additionally, the long-wavelength fluorescence at 715 nm in *P. tricornutum* that was retained in thylakoids isolated by using high-osmotic and high-ionic strength medium (Chrystal and Larkum, 1988) could be attributable to F710 emitted from FCP (Lhcf15) functionally connected to PSII (Herbstová et al., 2015) since F710 disappeared quickly *in vitro* as described above. This instability of F710 could be reworded by the dissociation of functional coupling between PSII core complexes and FCP responsible for F710 as described above. It was also indicated that protein synthesis is necessary for the development of energy flow to F710 under weak red light (Fujita and Ohki, 2004). The changes in fluorescence spectra observed in this work also required time longer than 6 h (**Fig. 7**). Accordingly, FL715 should contain F710 component in addition to PSI fluorescence. Then, the re-localization of antennas mentioned above includes placement/displacement of Lhcf15 with PSII in case of *P. tricornutum*.

### Distribution of Excitation Energy During Growth Irradiance Acclimation

State transitions are the mechanism that enables plants to redistribute excitation energy between PSI and PSII (Murata, 1969; Takahashi et al., 2006). State transitions are rapid acclimation process (occurring within ∼10 min) to respond to short-term changes in irradiance or light-quality and are triggered by the reduction of plastoquinone pool (Takahashi et al., 2006). As a result, the peripheral antenna complexes, LHCII, are translocated between PSI and PSII core complexes. The results obtained here suggest a similar regulation system in diatoms, in that FCPs are re-localized between PSI and PSII according to differences in growth irradiance. However, it is reported that state transitions are absent from diatoms (**Fig. 7**) (Owens, 1986). The changes that we observed occurred during less than two-generations; thus, it represents ‘state conversion’, which is an antenna rearrangement during the long-term acclimation process in diatoms, rather than ‘state transitions’. State transitions in green plants are the short-term acclimation process shuttling LHCII between PSII-LHCII and PSI-LHCI, consisting of the reactions of disassembly of LHCII from PSII-LHCII and assembly of phosphorylated LHCII to PSI-LHCI as state 2 and the reverse reactions. The assessment of time range necessary for state conversion is the future work since we could not determine during 6 h light-exposure experiment in this work (**Fig. 7**).

A schematic model of the regulation process based on our results is shown in **Fig. 10**. Some FCPs (brown) are tightly bound to PSI and PSII (Ikeda et al., 2008; Nagao et al., 2022, 2020, 2019, 2010; Xu et al., 2020); however, a large amount of FCPs (orange) is mobile and they could shuttle and balance excitation between the two photosystems (**Fig. 10**). The mobile FCPs might correspond to M- or L-type LHCII-trimers in green plants (Dekker and Boekema, 2005). Among the six subfamilies of FCPs, Lhcq proteins compose peripheral part of FCPI associated with PSI (Xu et al., 2020). This peripheral part of FCPI can be detached from the complex (Nagao et al., 2020) and, interestingly, the number of genes encoding Lhcq differs among diatom species. Therefore, it is reasonably assumed that the structure and composition of outer periphery are different under different growth conditions or among diatom species.

**Fig. 10.**
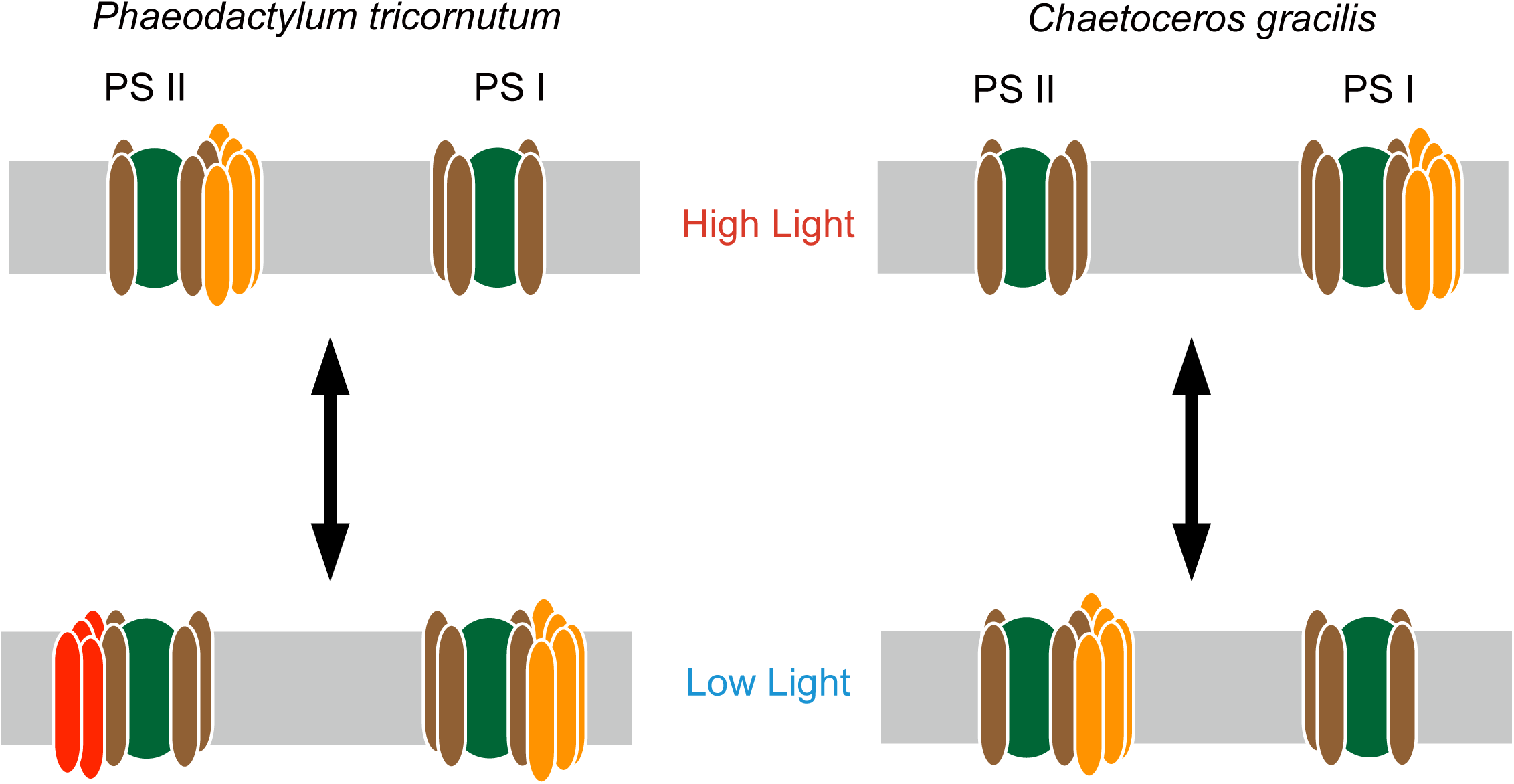
Schematic model of state conversion in two marine diatoms. PSI and PSII core complexes (green), bound FCPs (dark brown), mobile FCPs (orange), inducible FCP (F710) and thylakoid membrane (gray band). Changes in the proportions of PSI and PSII in cells under different growth irradiance are not considered. In *P. tricornutum* cells, most mobile FCPs are bound to PSII core under high light conditions but are translocated to PSI core under low light conditions. In contrast, in *C. gracilis* cells, most mobile FCPs are bound to PSI core under high-light conditions but are translocated to PSII core under low-light conditions. The mobile FCPs are not necessarily the same between PSI-FCPI-bound ones and PSII-FCPII-bound ones. In *P. tricornutum*, under low irradiances, F710 is induced to associate with PSII-FCPII. These growth irradiance-dependent re-arrangements of FCPs between PSI and PSII are designated as state conversion. Although state transitions are induced by the redox state of the plastoquinone pool and are irreversible in short periods, the translocations of the state conversion are slow and irreversible for at least six hours.

Lhcfs in diatoms form diverse oligomeric states, at least dimer, trimer, tetramer, and pentamer, which are classified into rigid or mobile FCPs collectively associated with PSII tightly or loosely. The different oligomeric states were originally found in *Cyclotella meneghiniana* (Büchel, 2003). In *T. pseudonana*, closely related to *C. meneghiniana* which belongs to the order Thalassiosirales as *T. pseudonana*, FCP trimers and FCP dimers are present. FCP trimers are homotrimers composed of TpLhcf8/9 (Zhou et al., 2024), which were not identified in the cryo-EM structure of PSII-FCPII (Feng et al., 2023; Nagao et al., 2022) although these FCP trimers are thought to associate with the outer periphery of PSII (Zhou et al., 2024). TpLhcf7 forms a homodimer associating with PSII at CP47 side in the cryo-EM structure of PSII-FCPII (Feng et al., 2023). TpLhcf5 and TpLhcf6 are also detected in the band of dimers on native PAGE (Zhou et al., 2024) and assigned in the cryo-EM structure of PSII-FCPII, forming a heterodimer unit (Feng et al., 2023). TpLhcf1/2 compose FCP homodimers (Zhou et al., 2024), which are not assigned in the cryo-EM structure of PSII-FCPII. TpLhcf1–7 and TpLhcf11 were detected in the FCPII band on native PAGE (Grouneva et al., 2011; Nagao et al., 2013), and additionally, they were also detected in the PSII-FCPII fraction on native PAGE (Calvaruso et al., 2020). TpLhcr17 (formerly annotated as TpLhca2 assuming to be the member of green-lineage Lhca subfamily but re-annotated as the red-lineage Lhcr subfamily member), TpLhcf5–7, and TpLhcx6_1 were assigned in cryo-EM structure of PSII-FCPII (Feng et al., 2023). Accordingly, note that not all FCPII were assigned in the cryo-EM structure of *T. pseudonana* PSII-FCPII, which means that the *in vivo* functional PSII-FCPII accompanies loosely associated FCPII, maybe including mobile trimers and dimers. In addition, there are different affinities or binding orders; trimers are not associated with PSII both in native PAGE and cryo-EM structure, whereas TpLhcf1/2 homodimers were detected in the PSII-FCPII band in native PAGE. These facts indicate that the FCP trimer is more easily detached during the isolation process than TpLhcf1/2 dimers, suggesting TpLhcf1/2 homodimers bind to PSII-FCPII in a more physically rigid or closer manner. Therefore, the mobile FCPs are different protein species from those that bind tightly to PSI and PSII.

In contrast, *C. gracilis* has FCP dimers, tetramers, and pentamers. *C. gracilis* PSII-FCPII contains tetramers with three CgLhcf1s and one CgLhcf5, and pentamers consisting of one unit of tetramer in a rotationally symmetric manner with two CgLhcf1s, one CgLhcf6, and one CgLhcf7, accompanying CgLhcf13 at CgLhcf7 side. FCP dimers are not found in cryo-EM structures of *C. gracilis* PSII-FCPII nor in that of PSI-FCPI (Nagao et al., 2022, 2020; Xu et al., 2020). It is estimated that the dimers in *C. gracilis* consist of CgLhcf2 and CgLhcf12, originally annotated as FCP-3/4 sharing the HGRIAQLAFLGN sequence identified by amino-acid sequencing (Ishihara et al., 2015). This estimation was confirmed by isolation of the dimers, resulting in heterodimers of CgLhcf2 and CgLhcf12 (Zhou et al. 2024). CgLhcf2–CgLhcf12 dimers should be the mobile FCPs in *C. gracilis*, in which it harbors ∼60% of cellular Chl *a* (Ishihara et al., 2015), and would contribute to rebalancing the excitation between two photosystems. The information regarding oligomerization of FCPs in *P. tricornutum* is very limited. While most of the FCPs in *P. tricornutum* are found to be dimers on clear native PAGE (Nagao et al., 2013), only the structure of PtLhcf4 homodimer was reported, and higher order assembly of *P. tricornutum* FCPII is not well known except F710 described below (Herbstová et al., 2017). The cryo-EM structures of PSI-FCPI and PSII-FCPII are not yet available. Therefore, currently, we do not have enough information to presume mobile FCPs in *P. tricornutum*.

There should also be a group of FCP responsible for the major component of non-photochemical fluorescence quenching coupled with the xanthophyll cycle to protect photosystems from incidental high light in diatoms (Kashino and Kudoh, 2003; Olaizola and Yamamoto, 1994) (see (Goss and Lepetit, 2015) for review). An FCP protein with this function was identified in the centric diatom *C. meneghiniana* (Beer et al., 2006) and named as Lhcx. Lhcx proteins were called as LI818-like proteins derived from the former name (LI818) of one type LHC (Lhcsr) in green algae (Zhu and Green, 2010), which play critical roles in the energy dependent quenching (qE) capacity (Peers et al., 2009). Diatoms have several Lhcx proteins serving for qE as well (see (Goss and Lepetit, 2015) for review). In recent years, studies using knock-down, knock-out, and overexpressing lines of the Lhcx proteins revealed the characteristics and contribution to qE capacity of each Lhcx proteins reflecting environmental signals and nutrient availability (Bailleul et al., 2010; Buck et al., 2022, 2021, 2019; Giovagnetti et al., 2022; Hao et al., 2018; Taddei et al., 2018). Characterizations of FCPs from *P. tricornutum* grown under low and high irradiance (40 and 140 µmol photons·m^−2^·s^−1^ in (Gundermann et al., 2013); 30 and 300 µmol photons·m^−2^·s^−1^ in (Nagao et al., 2021)) were reported. These FCPs contained only Lhcf and Lhc-like proteins (Gundermann et al., 2013) or only Lhcf, Lhcr and Lhcq proteins (Nagao et al., 2021). Also, the FCP-trimer complex purified from transgenic *P. tricornutum* expressing His-tagged FcpA consisted of only two types of FCPs (Joshi-Deo et al., 2010). These findings are consistent with the previous reports in *C. gracilis* (Ishihara et al., 2015). Therefore, the mobile FCPs likely contain little, if any, Lhcx proteins in *P. tricornutum*, especially when grown under high light. The expression mechanism and physiological functions of Lhcx are now better clarified by studies using genome editing for *P. tricornutum* (Concordet and Haeussler, 2018; Zhang et al., 2024).

The F710 fluorescence emission in *P. tricornutum* observed in our experiments may originate from low-energy, long-wavelength chlorophyll species within the PtLhcf15 antenna complex (Herbstová et al., 2015), which function as energetic sinks under 77K conditions. Although the PtLhcf15 oligomer works as peripheral FCPII, the excitation energy transfer from PtLhcf15 to PSII is an uphill process. However, at 77K, thermal activation becomes negligible, effectively blocking this uphill energy transfer pathway and resulting in the pronounced F710 fluorescence emission observed. The low-energy chlorophyll sites in PtLhcf15 can also act as energy sinks that can accept excitation energy from PSII-FCPII complexes through downhill energy transfer. The observed changes in FL715 intensity following freeze-thaw cycles can be attributed to alterations in the relative contributions of these energy flow pathways: direct energy capture within PtLhcf15 leading to F710 emission versus energy spillover from PSII-FCPII to the F710-emitting states, potentially due to disruption of the correct coordination of chromophores within the antenna-reaction center supercomplex.

Changes in the fluorescence spectra of *P. tricornutum* similar to those reported here were previously documented by Herbstová et al. (Herbstová et al., 2015). In their study, fluorescence at 681 nm was dominant in daylight-grown cells, while fluorescence at 712 nm was dominant in cells grown under red-enhanced ambient light, with the emitter denoted as F710 and attributed to an oligomer of PtLhcf15 (Herbstová et al., 2017, 2015). Their cells were cultivated under red-enhanced light of ∼20 µmol photons·m^−2^·s^−1^ using an incandescent bulb, conditions similar to our low-light experimental setup. Herbstová and colleagues interpreted this phenomenon as chromatic adaptation, assuming that red light becomes dominant underwater in natural environments (Herbstová et al., 2017, 2015). However, the opposite is true—red light is substantially reduced while blue light becomes dominant in the water column (Falkowski and Raven, 2007; Kirk, 1994). Based on our findings, both sets of results can be more appropriately interpreted through our state conversion model. In *P. tricornutum*, the expression of PtLhcf15 enhances the antenna size of PSII under dimmer light conditions, with partial supplementation of far-red-light absorption capacity.

The interaction between PSII-FCPII and PtLhcf15 appears to be inherently weak, such that this association is readily disrupted following freeze-thaw treatment (this work) or during thylakoid isolation procedures (Brown, 1967; Fujita and Ohki, 2004; Herbstová et al., 2015; Sugahara et al., 1971). This disruption results in decreased energy transfer efficiency and alters the coordination of chromophores within the supercomplex, leading to the observed changes in F710 (or FL715) fluorescence emission. Therefore, it is more reasonable to interpret the results reported by Herbstová et al. (Herbstova et al., 2015b) in the context of light-dependent state conversion rather than chromatic adaptation to spectral light quality. It has not been clarified yet whether *C.gracilis* have Lhc with low-energy, long-wavelength chlorophyll species; even if it does, the patterns of accumulation and gene expression are expected to differ from those of PtLhcf15.

## CONCLUSION

A change in the proportions of PSI and PSII under different irradiances can occur to some extent, as reported elsewhere (Falkowski and Owens, 1980; Falkowski et al., 1981; Smith and Melis, 1988; Strzepek and Harrison, 2004). However, our results indicate that the mobile and inducible FCPs balance the photochemical excitation between PSI and PSII for efficient photosynthesis of diatoms under the wide range of growth irradiances in their habitat, from the surface to the depths of the ocean.

In our study, the two diatoms *C. gracilis* and *P. tricornutum* showed apparently opposite changes in the proportions of light-energy delivery to PSII and PSI. This opposite change in the proportions of photosystems in diatoms, such as *T. costatum*, *T. oceania*, *T. weissflogii*, *C. fusiformis*, has been reported in other studies (Falkowski and Owens, 1980; Smith and Melis, 1988; Strzepek and Harrison, 2004). This indicates the wide variety of acclimation strategies of diatoms that inhabit diverse natural environments to changes in environmental conditions. To fully understand the acclimation strategy of diatoms in terms of photosynthesis, it is important to analyze many aspects of photosynthesis including linear and cyclic electron flow, state conversion, carbon assimilation, and light protection in the context of their natural environment.

## MATERIALS AND METHODS

### Diatom Strains and Growth Conditions

The centric marine diatom *Chaetoceros gracilis* Schütt (UTEX LB 2658) and the pennate marine diatom *Phaeodactylum tricornutum* Böhlin (UTEX 642) were grown photoautotrophically in F/2 (Guillard and Ryther, 1962) or Provasoli’s (Provasoli, 1968) artificial sea water medium under continuous light, which was bubbled with air at 20°C (Ikeda et al., 2008) for approximately two generations before analysis. Cell density was measured by determining optical density at 750 nm (OD750) with an MPS-2000 spectrophotometer (Shimadzu, Kyoto, Japan). For all experiments, the initial cell density (OD750) was ∼0.02. Growth irradiance was controlled by changing the distance from the light source (two 150 W halogen lamps; TRAD HL-150, Sankyo, Osaka, Japan) to achieve 3, 15, 70, or 320 µmol photons·m^−2^·s^−1^. Ultraviolet radiation that could affect the fluorescence properties of diatoms (Goedheer, 1973) and heat from halogen lamps was dissipated using a 5-cm-thick water filter in square glass bottles. Irradiances were determined using a quantum photometer (Licor, Inc., Lincoln, NE, USA). The cells were harvested by centrifugation at 25,000 *g* for 5 min at 10°C, washed, and re-suspended in a medium A (0.5 M sucrose, 20 mM MES-NaOH (pH 6.7), 10 mM NaCl, and 5 mM MgCl_2_).

### Purification of Photosystems I and II from *Phaeodactylum tricornutum*

*P. tricornutum* cells were grown photoautotrophically in 7.5 L artificial sea water (Marine Art SF-1; Tomita Pharmaceutical, Tokushima, Japan) supplemented with Daigo’s IMK medium for marine microalgae (Wako, Osaka, Japan). The cultures were aerated with air, and grown at 20°C for 15 days in a square 9-L polycarbonate bottle (Nalge Nunc, Rochester, NY). Continuous light was supplied by one 40 W light bulb for 6 d and then by two 40 W light bulbs for the remaining days. Cells were collected from late logarithmic-phase cultures by centrifugation (7,800 *g*, 8 min) and resuspended in medium B (10 mM MgCl_2_, 5 mM CaCl_2_, 50 mM MES-NaOH (pH6.5), 5% glycerol, and 1 M betaine). Cells equivalent to 1 mg Chl *a*/mL were broken with the same volume of glass beads (φ0.105~0.125 mm; BZ-01, AS ONE Corp, Osaka Japan) at 4°C by 20 cycles of 5-s breaking and 2-min cooling in the presence of DNase I (0.5 µg/ml, Sigma, St Louis, MO, USA) and a protease inhibitor mixture (250 µL/100 mL cell suspension, Sigma). Thylakoid membranes were collected by centrifuging the broken cells at 27,000 *g* for 15 min and then resuspended in medium B. To purify PSI and PSII, thylakoid membranes were solubilized in 3.0% *n*-dodecyl-β-D-maltoside (DDM, Anatrace, Maumee, OH) for 15 min on ice, to a final concentration of 1.0 mg Chl *a*/mL. The resulting extracts were fractionated by sucrose density gradient ultracentrifugation at 108,000 *g* for 18.5 h at 4°C. The stepwise sucrose density gradient comprised 4-mL each of 0.3, 0.5, 0.7, 1.0 and 1.2 M sucrose in a medium consisting of 50 mM MES-NaOH (pH 6.5), 10 mM MgCl_2_, 5 mM CaCl_2_, and 0.04% DDM. The resulting two green bands were collected, and the proteins were precipitated by adding the same volume of 25% polyethylene glycol 2000 monomethyl ether (Fluka, Steinheim, Germany) and then resuspended in medium B supplemented with 0.04% DDM.

### Polypeptide Analysis

Polypeptides were separated by sodium dodecyl sulfate-polyacrylamide gel electrophoresis (SDS-PAGE) (Kashino et al., 2001) on gels containing 18% acrylamide and 6 M urea. Specific antibodies were used to detect reaction center proteins of PSI (PsaA/B) (Kashino et al., 1990) and PSII (PsbA) (anti-PsbA global; Agri Sera, Vännäs, Sweden). Immuno-decorated bands were detected by enhanced chemiluminescence with a LAS-4000 mini (GE Healthcare, Buckinghamshire, England) using WestFemto reagents (Pierce, Rockford, IL, USA).

### Spectroscopic Analysis

Steady-state fluorescence emission spectra at 77K were measured using an H-20 UV monochromator (Jobin Yvon, Cedex, France) as described previously (Inoue-Kashino et al., 2005) or a FluoroMax-4 spectrofluorometer (Horiba, Tokyo, Japan). When using the H-20 UV monochromator, chlorophylls were excited by blue light, which was supplied by a halogen lamp and passed through a Corning 4-96 optical band-pass filter (Corning, Corning, NY, USA). The fluorescence spectra were plotted against wavenumber (inverse of the wavelength) and fitted to two Lorentzian components using KaleidaGraph (Synergy Software, Reading, PA, USA) with a Levenberg–Marquardt regression algorithm when necessary.

### Measurements of light-induced oxidation of P700

Light-induced absorption changes of P700, the reaction center pigment in PSI, in cells were monitored with a Joliot type-spectrometer JTS-10 (BioLogic, Claix, France). The reaction mixture contained 5 µg Chl *a*/mL in the presence of 10 µM 3-(3,4- dichlorophenyl)-1,1-dimethylurea (DCMU), 20 µM 2,5-dibromo-3-methyl-6-isopropyl-p-benzoquinone (DBMIB), 2.5 mM methylviologen, and 5 mM sodium ascorbate. Actinic light (AL) (84 µmol photons·m^−2^·s^−1^ at 630 nm) oxidized P700, and the absorption changes were measured at 705 nm.

### Pigment Analysis

Photosynthetic pigments were quantified by reversed-phase HPLC as described previously (Kashino and Kudoh, 2003), after extraction in *N,N*-dimethylformamide (Furuya et al., 1998; Hashihama et al., 2010). Standard pigments were purchased from the VKI Water Quality Institute (Hørsholm, Denmark).

## DATA AVAILABILITY STATEMENT

Data sharing is not applicable to this article as all new created data is already contained within this article.

## FUNDING INOFRMATION

This work was supported by JST-ALCA (JPMJAL1105, JPMJAL1608 to Y.K. and K.I.), MEXT JSPS KAKENHI (25740054 to N. I.-K.; 18054028, 18GS0318 to Y.K.; 20H03116 to K.I. and Y.K.; 20K06234 to S.A.; 22KJ2017 to M.K.; 16H06554, 21H02510 to YT), grants from the Hyogo Science and Technology Association (20I107 to Y.K.), Sasakawa Scientific Research Grants from The Japan Science Society (21-417 to N. I.-K. and 24-444 to T.I.), Akira Yoshino Research Grant of The Chemical Society of Japan (K.I.), and the National Institute of Polar Research (Y.K.).

## ACKNOWLEDGENTS

We thank Ms. Chie Minami for her technical assistance.

## AUTHOR CONTRIBUTIONS

Y.K. and S.K. conceived the original screening and research plans; K.I., K.S. and Y.T. supervised the experiments; N.I.-K. performed most of the experiments and analyzed the data; S.A. and T.I. provided technical assistance to N.I.-K.; S.A. and K.F. performed initial screening experiments; Y.K. conceived the project, designed the experiments and wrote the article with contributions of all the authors; M.K., S.A., K.I. and Y.T. complemented the writing.

## Abbreviations

Chl,: chlorophyll;
PSI and PSII,: photosystem I and II, respectively;

## REFERENCES

1. Arsalane, W., Rousseau, B., and Duval, J.-C. (1994) Influence of the pool size of the xanthophyll cycle on the effects of light stress in a diatom: Competition between photoprotection and photoinhibition. Photochem Photobiol 60: 237–243.

2. Bailleul, B., Rogato, A., De Martino, A., Coesel, S., Cardol, P., Bowler, C., et al. (2010) An atypical member of the light-harvesting complex stress-related protein family modulates diatom responses to light. Proc Natl Acad Sci USA 107: 18214–18219.

3. Ban, A., Aikawa, S., Hattori, H., Sasaki, H., Sampei, M., Kudoh, S., et al. (2006) Comparative analysis of photosynthetic properties in ice algae and phytoplankton inhabiting Franklin Bay, the Canadian Arctic, with those in mesophilic diatoms during CASES 03-04. Polar Biosci 19: 11–28.

4. Beer, A., Gundermann, K., Beckmann, J., and Buchel, C. (2006) Subunit composition and pigmentation of fucoxanthin-chlorophyll proteins in diatoms: evidence for a subunit involved in diadinoxanthin and diatoxanthin binding. Biochemistry 45: 13046–13053.

5. Brown, J.S. (1967) Fluorometric evidence for the participation of chlorophyll *a*-695 in system 2 of photosynthesis. Biochim Biophys Acta 143: 391–398.

6. Büchel, C. (2003) Fucoxanthin-chlorophyll proteins in diatoms: 18 and 19 kDa subunits assemble into different oligomeric states. Biochemistry 42: 13027–13034.

7. Buck, J.M., Kroth, P.G., and Lepetit, B. (2021) Identification of sequence motifs in Lhcx proteins that confer qE-based photoprotection in the diatom *Phaeodactylum tricornutum*. Plant Journal 108: 1721–1734.

8. Buck, J.M., Sherman, J., Bártulos, C.R., Serif, M., Halder, M., Henkel, J., et al. (2019) Lhcx proteins provide photoprotection via thermal dissipation of absorbed light in the diatom *Phaeodactylum tricornutum*. Nat Commun 10: 4167.

9. Buck, J.M., Wünsch, M., Schober, A.F., Kroth, P.G., and Lepetit, B. (2022) Impact of Lhcx2 on Acclimation to Low Iron Conditions in the Diatom *Phaeodactylum tricornutum*. Front Plant Sci 13: 841058.

10. Calvaruso, C., Rokka, A., Aro, E.M., and Buchel, C. (2020) Specific Lhc Proteins Are Bound to PSI or PSII Supercomplexes in the Diatom *Thalassiosira pseudonana*. Plant Physiol 183: 67–79.

11. Chrystal, J., and Larkum, A.W.D. (1988) Preservation of long-wavelength fluorescence in the isolated thylakoids of two phytoplanktonic algae at 77 K. Biochim Biophys Acta 932: 189–194.

12. Concordet, J.-P., and Haeussler, M. (2018) CRISPOR: intuitive guide selection for CRISPR/Cas9 genome editing experiments and screens. Nucleic Acids Res 46: W242–W245.

13. Dekker, J.P., and Boekema, E.J. (2005) Supramolecular organization of thylakoid membrane proteins in green plants. Biochim Biophys Acta 1706: 12–39.

14. Dilworth, M.F., and Gantt, E. (1981) Phycobilisome-thylakoid topography on photosynthetically active vesicles of *Porphyridium cruentum*. Plant Physiol 67: 608–612.

15. van Dorssen, R.J., Plijter, J.J., Dekker, J.P., den Ouden, A., Amesz, J., and van Gorkom, H.J. (1987) Spectroscopic properties of chloroplast grana membranes and of the core of Photosystem II. Biochim Biophys Acta 890: 134–143.

16. Falciatore, A., Jaubert, M., Bouly, J.P., Bailleul, B., and Mock, T. (2020) Diatom molecular research comes of age: Model species for studying phytoplankton biology and diversity. Plant Cell 32:547–572.

17. Falkowski, P., and Owens, T.G. (1980) Light-shade adaptation: Two strategies in marine phytoplankton. Plant Physiol 66: 592–595.

18. Falkowski, P.G., Owens, T.G., Ley, A.C., and Mauzerall, D.C. (1981) Effects of growth irradiance levels on the ratio of reaction centers in two species of marine phytoplankton. Plant Physiol 68: 969–973.

19. Falkowski, P.G., and Raven, J.A. (2007) Aquatic Photosynthesis, 2nd ed. Princeton University Press, New Jersey

20. Feng, Y., Li, Z., Li, X., Shen, L., Liu, X., Zhou, C., et al. (2023) Structure of a diatom photosystem II supercomplex containing a member of Lhcx family and dimeric FCPII. Sci Adv 9; eadi8446.

21. Feng, Y., Li, Z., Yang, Y., Shen, L., Li, X., Liu, X., et al. (2025) Structures of PSI–FCPI from *Thalassiosira pseudonana* grown under high light provide evidence for convergent evolution and light-adaptive strategies in diatom FCPIs. J Integr Plant Biol 67: 949–966.

22. Fujita, Y., and Ohki, K. (2004) On the 710 nm fluorescence emitted by the diatom *Phaeodactylum tricornutum* at room temperature. Plant Cell Physiol 45: 392–397.

23. Furuya, K., Hayashi, M., and Yabushita, Y. (1998) HPLC determination of phytoplankton pigments using N,N-dimethylformamide. J Oceanogr 54: 199–203.

24. Giovagnetti, V., Jaubert, M., Shukla, M.K., Ungerer, P., Bouly, J.P., Falciatore, A., et al. (2022) Biochemical and molecular properties of LHCX1, the essential regulator of dynamic photoprotection in diatoms. Plant Physiol 188: 509–525.

25. Goedheer, J.C. (1973) Chlorophyll *a* forms in *Phaeodactylum tricornutum*: comparison with other diatoms and brown algae. Biochim Biophys Acta 314: 191–201.

26. Goss, R., and Lepetit, B. (2015) Biodiversity of NPQ. J Plant Physiol 172: 13–32.

27. Grouneva, I., Rokka, A., and Aro, E.M. (2011) The thylakoid membrane proteome of two marine diatoms outlines both diatom-specific and species-specific features of the photosynthetic machinery. J Proteome Res 10: 5338–5353.

28. Guillard, R.R., and Ryther, J.H. (1962) Studies of marine planktonic diatoms. I. *Cyclotella nana* Hustedt, and *Detonula confervacea* (cleve) Gran. Can J Microbiol 8: 229–239.

29. Gundermann, K., Schmidt, M., Weisheit, W., Mittag, M., and Büchel, C. (2013) Identification of several sub-populations in the pool of light harvesting proteins in the pennate diatom *Phaeodactylum tricornutum*. Biochim Biophys Acta 1827: 303–310.

30. Hao, T. Bin, Jiang, T., Dong, H.P., Ou, L. jian, He, X., and Yang, Y. F. (2018) Light-harvesting protein Lhcx3 is essential for high light acclimation of *Phaeodactylum tricornutum* AMB Express 8: 174.

31. Hashihama, F., Umeda, H., Hamada, C., Kudoh, S., Hirawake, T., Satoh, K., et al. (2010) Light acclimation states of phytoplankton in the Southern Ocean, determined using photosynthetic pigment distribution. Mar Biol 157: 2263–2278.

32. Herbstová, M., Bína, D., Kaňa, R., Vácha, F., and Litvín, R. (2017) Red-light phenotype in a marine diatom involves a specialized oligomeric red-shifted antenna and altered cell morphology. Sci Rep 7: 11976.

33. Herbstová, M., Bina, D., Konik, P., Gardian, Z., Vacha, F., and Litvin, R. (2015) Molecular basis of chromatic adaptation in pennate diatom *Phaeodactylum tricornutum*. Biochim Biophys Acta 1847: 534–543.

34. Hihara, Y., and Sonoike, K. (2001) Regulation, Inhibition and Protection of Photosystem I. In Regulation of Photosynthesis. Edited by Aro, E.M. and Andersson, B. pp. 507–531 Kluwer Academic Press, Dordrecht, The Netherlands.

35. Ifuku, K., Yan, D., Miyahara, M., Inoue-Kashino, N., Yamamoto, Y.Y., and Kashino, Y. (2015) A stable and efficient nuclear transformation system for the diatom *Chaetoceros gracilis*. Photosynth Res 123: 203–211.

36. Ikeda, Y., Komura, M., Watanabe, M., Minami, C., Koike, H., Itoh, S., et al. (2008) Photosystem I complexes associated with fucoxanthin-chlorophyll-binding proteins from a marine centric diatom, *Chaetoceros gracilis*. Biochim Biophys Acta 1777: 351–361.

37. Ikeda, Y., Satoh, K., and Kashino, Y. (2005) Characterization of photosystem I complexes purified from a diatom, *Chaetoceros gracilis*. In Photosynthesis: Fundamental Aspects to Global Perspectives. Edited by van der Est, A. and Bruce, D. pp. 38–40 Alliance Communications Group, Kansas.

38. Inoue-Kashino, N., Kashino, Y., Satoh, K., Terashima, I., and Pakrasi, H.B. (2005) PsbU provides a stable architecture for the oxygen-evolving system in cyanobacterial photosystem II. Biochemistry 44: 12214–12228.

39. Ishihara, T., Ifuku, K., Yamashita, E., Fukunaga, Y., Nishino, Y., Miyazawa, A., et al. (2015) Utilization of light by fucoxanthin-chlorophyll-binding protein in a marine centric diatom, *Chaetoceros gracilis*. Photosynth Res 126: 437–447.

40. Itoh, S., Sugiura, K., and Govindjee (2004) Fluorescence of photosystem I. In Chlorophyll a Fluorescence: A Signature of Photosynthesis. Edited by Papageorgiou, G.C. and Govindjee. pp. 231–250 Springer, Dordrech, The Netherlands.

41. Joshi-Deo, J., Schmidt, M., Gruber, A., Weisheit, W., Mittag, M., Kroth, P.G., et al. (2010) Characterization of a trimeric light-harvesting complex in the diatom *Phaeodactylum tricornutum* built of FcpA and FcpE proteins. J Exp Bot 61: 3079–3087.

42. Kashino, Y., Enami, I., Satoh, K., and Katoh, S. (1990) Immunological cross-reactivity among corresponding proteins of photosystems I and II from widely divergent photosynthetic organisms. Plant Cell Physiol 31: 479–488.

43. Kashino, Y., Koike, H., and Satoh, K. (2001) An improved sodium dodecyl sulfate-polyacrylamide gel electrophoresis system for the analysis of membrane protein complexes. Electrophoresis 22: 1004–1007.

44. Kashino, Y., and Kudoh, S. (2003) Concerted response of xanthophyll-cycle pigments in a marine diatom, *Chaetoceros gracilis*, to the sifts of light condition. Phycol Res 51: 168–172.

45. Kashino, Y., Lauber, W.M., Carroll, J.A., Wang, Q., Whitmarsh, J., Satoh, K., et al. (2002) Proteomic analysis of a highly active photosystem II preparation from the cyanobacterium Synechocystis sp. PCC 6803 reveals the presence of novel polypeptides. Biochemistry 41: 8004–8012.

46. Katoh, T., and Gantt, E. (1979) Photosynthetic vesicles with bound phycobilisomes from *Anabaena variabilis*. Biochim Biophys Acta 546: 383–393.

47. Kirk, J.T.O. (1994) Light & Photosynthesis in Aquatic Ecosystems, 2nd Edition. ed. Cambridge University Press, Cambridge.

48. Kumazawa, M., Nishide, H., Nagao, R., Inoue-Kashino, N., Shen, J.R., Nakano, T., et al. (2022) Molecular phylogeny of fucoxanthin-chlorophyll *a*/*c* proteins from *Chaetoceros gracilis* and Lhcq/Lhcf diversity. Physiol Plant 174: e13598.

49. Kura-Hotta, M., Satoh, K., and Katoh, S. (1986) Functional linkage between phycobilisome and reaction center in two phycobilisome oxygen-evolving photosystem II preparations isolated from the thermophilic cyanobacterium *Synechococcus* sp. Arch Biochem Biophys 249: 1–7.

50. De la Torre, W.R., and Burkey, K.O. (1990) Acclimation of barley to changes in light intensity: chlorophyll organization. Photosynth Res 24: 117–125.

51. Lamb, J.J., Røkke, G., and Hohmann-Marriott, M.F. (2018) Chlorophyll fluorescence emission spectroscopy of oxygenic organisms at 77 K. Photosynthetica 56: 105–124.

52. Lepetit, B., Volke, D., Szabo, M., Hoffmann, R., Garab, G., Wilhelm, C., et al. (2007) Spectroscopic and molecular characterization of the oligomeric antenna of the diatom *Phaeodactylum tricornutum*. Biochemistry 46: 9813–9822.

53. Macpherson, A.N., and Hiller, R.G. (2003) Light-Harvesting Systems in Chlorophyll c-Containing Alage. In Light-Harvesting Antennas in Photoysnthesis. Edited by Green, B.R. and Parson, W.W. pp. 323–352 Kluwer Academic Publishers, Dordrecht, The Netherlands.

54. Mimuro, M. (2004) Photon Capture, Exciton Migration and Trapping and Fluorescence Emission in Cyanobacteria and Red Alagae. In Chlorophyll a Fluorescence: A Signature of Photosynthesis. Edited by Papageorgiou, G.C. and Govindjee. pp. 173–195 Springer, Dordrech, The Netherlands.

55. Miyahara, M., Aoi, M., Inoue-Kashino, N., Kashino, Y., and Ifuku, K. (2013) Highly efficient transformation of the diatom *Phaeodactylum tricornutum* by multi-pulse electroporation. Biosci Biotechnol Biochem 77: 874–876.

56. Murakami, A. (1997) Quantitative analysis of 77K fluorescence emission spectra in *Synechocystis sp.* PCC 6714 and *Chlamydomonas reinhardtii* with variable PS I/PS II stoichiometries. Photosynth Res 53: 141–148.

57. Murata, N. (1969) Control of excitation transfer in photosynthesis. I. Light-induced change of chlorophyll *a* fluoresence in *Porphyridium cruentum*. Biochim Biophys Acta 172: 242–251.

58. Murata, N., Nishimura, M., and Takamiya, A. (1966) Fluorescence of chlorophyll in photosynthetic systems. III. Emission and action spectra of fluorescence-three emission bands of chlorophyll a and the energy transfer between two pigment systems. Biochim Biophys Acta 126: 234–243.

59. Murata, N., and Satoh, K. (1986) Absorption and Fluorescence Emission by Intact Cells, Chloroplasts, and Chlorophyll-Protein Complexes. In Light Emission by Plants and Bacteria. Edited by Govindjee, Amesz, J., and Ford, D.C. pp. 137–159 Academic Press, USA, Orlando.

60. Nagao, R., Ishii, A., Tada, O., Suzuki, T., Dohmae, N., Okumura, A., et al. (2007) Isolation and characterization of oxygen-evolving thylakoid membranes and Photosystem II particles from a marine diatom *Chaetoceros gracilis*. Biochim Biophys Acta 1767: 1353–1362.

61. Nagao, R., Kato, K., Ifuku, K., Suzuki, T., Kumazawa, M., Uchiyama, I., et al. (2020) Structural basis for assembly and function of a diatom photosystem I-light-harvesting supercomplex. Nat Commun 11: 2481.

62. Nagao, R., Kato, K., Kumazawa, M., Ifuku, K., Yokono, M., Suzuki, T., et al. (2022) Structural basis for different types of hetero-tetrameric light-harvesting complexes in a diatom PSII-FCPII supercomplex. Nat Commun 13: 1764.

63. Nagao, R., Kato, K., Suzuki, T., Ifuku, K., Uchiyama, I., Kashino, Y., et al. (2019) Structural basis for energy harvesting and dissipation in a diatom PSII-FCPII supercomplex. Nat Plants 5: 890–901.

64. Nagao, R., Takahashi, S., Suzuki, T., Dohmae, N., Nakazato, K., and Tomo, T. (2013) Comparison of oligomeric states and polypeptide compositions of fucoxanthin chlorophyll *a*/*c*-binding protein complexes among various diatom species. Photosynth Res 117: 281–288.

65. Nagao, R., Tomo, T., Noguchi, E., Nakajima, S., Suzuki, T., Okumura, A., et al. (2010) Purification and characterization of a stable oxygen-evolving Photosystem II complex from a marine centric diatom, *Chaetoceros gracilis*. Biochim Biophys Acta 1797: 160–166.

66. Nagao, R., Yokono, M., Ueno, Y., Suzuki, T., Kumazawa, M., Kato, K.H., et al. (2021) Enhancement of excitation-energy quenching in fucoxanthin chlorophyll *a*/*c*-binding proteins isolated from a diatom *Phaeodactylum tricornutum* upon excess-light illumination. Biochim Biophys Acta Bioenerg 1862.

67. Nelson, D.M., Treguer, P., Brzezinski, M.A., Leynaert, A., and Queguiner, B. (1995) Production and dissolution of biogenic silica in the ocean: revised global estimates, comparison with regional data and relationship to biogenic sedimentation. Global Biogeochem Cycles 9: 359–372.

68. Olaizola, M., Laroche, J., Kolber, Z., and Falkowski, P.G. (1994) Non-photochemical fluorescence quenching and the diadinoxanthin cycle in a marine diatom. Photosynth Res 41: 357–370.

69. Olaizola, M., and Yamamoto, H.Y. (1994) Short-term response of the diadinoxanthin cycle and fluorescence yield to high irradiance in *Chaetoceros muelleri* (Bacillariophyceae). J Phycol 30: 606–612.

70. Owens, T.G. (1986) Light-harvesting function in the diatom *Phaeodactylum tricornutum*: II. Distribution of excitation energy between the photosystems. Plant Physiol 80: 739–746.

71. Peers, G., Truong, T.B., Ostendorf, E., Busch, A., Elrad, D., Grossman, A.G., et al. (2009) An ancient light-harvesting protein is critical for the regulation of algal photosynthesis. Nature 462: 518–521.

72. Pi, X., Zhao, S., Wang, W., Liu, D., Xu, C., Han, G., et al. (2019) The pigment-protein network of a diatom photosystem II-light-harvesting antenna supercomplex. Science 365: eaax4406.

73. Provasoli, L. (1968) Media and prospects for the cultivation of marine algae. In Cultures and Collections of Algae. Proc. U.S.-Japan Conf. Edited by Watanabe, A. and Hattori, A. pp. 63–75 Jap. Soc. Plant Physiol., Kyoto.

74. Ramachandra, T. V, Mahapatra, D.M., Karthick, B., and Gordon, R. (2009) Milking diatoms for sustainable energy: Biochemical engineering versus gasoline-secreting diatom solar panels. Ind Eng Chem Res 48: 8769–8788.

75. Rijgersberg, C.P., and Amesz, J. (1980) Fluorescence and energy transfer in phycobiliprotein-containing algae at low temperature. Biochim Biophys Acta 593: 261–271.

76. Scheer, H. (2003) The Pigments. In Light-Harvesting Antennas in Photosynthesis. Edited by Green, B.R. and Parson, W.W. pp. 29–81 Kluwer Academic Publishers, Dordrecht.

77. Shimura, S., and Fujita, Y. (1973) Some properties of the chlorophyll fluorescence of the diatom *Phaeodactylum tricornutum*. Plant Cell Physiol 14: 341–352.

78. Sims, P.A., Mann, D.G., and Medlin, L.K. (2006) Evolution of the diatoms: insights from fossil, biological and molecular data. Phycologia 4: 361–402.

79. Smith, B.M., and Melis, A. (1988) Photochemical apparatus organization in the diatom *Cylindrotheca fusiformis*: Photosystem stoichiometry and excitation distribution in cells grown under high and low irradiance. Plant Cell Physiol 29: 761–769.

80. Strzepek, R.F., and Harrison, P.J. (2004) Photosynthetic architecture differs in coastal and oceanic diatoms. Nature 431: 689–692.

81. Sugahara, K., Murata, N., and Takamiya, A. (1971) Fluorescence of chlorophyll in brown algae and diatoms. Plant Cell Physiol 12: 377–385.

82. Taddei, L., Chukhutsina, V.U., Lepetit, B., Stella, G.R., Bassi, R., van Amerongen, H., et al. (2018) Dynamic changes between two LHCX-related energy quenching sites control diatom photoacclimation. Plant Physiol 177.

83. Takahashi, H., Iwai, M., Takahashi, Y., and Minagawa, J. (2006) Identification of the mobile light-harvesting complex II polypeptides for state transitions in *Chlamydomonas reinhardtii*. Proc Natl Acad Sci USA 103: 477–482.

84. Tokushima, H., Inoue-Kashino, N., Nakazato, Y., Masuda, A., Ifuku, K., and Kashino, Y. (2016) Advantageous characteristics of the diatom *Chaetoceros gracilis* as a sustainable biofuel producer. Biotechnol Biofuels 9: 235.

85. Vinayak, V., Manoylov, K.M., Gateau, H., Blanckaert, V., Herault, J., Pencreac’h, G., et al. (2015) Diatom milking: a review and new approaches. Mar Drugs 13: 2629–2665.

86. Xu, C., Pi, X., Huang, Y., Han, G., Chen, X., Qin, X., et al. (2020) Structural basis for energy transfer in a huge diatom PSI-FCPI supercomplex. Nat Commun 11: 5081.

87. Zaslavskaia, L.A., Lippmeier, J.C., Shih, C., Ehrhardt, D., Grossman, A.R., and Apt, K.E. (2001) Trophic conversion of an obligate photoautotrophic organism through metabolic engineering. Science 292: 2073–2075.

88. Zhang, H., Xiong, X., Guo, K., Zheng, M., Cao, T., Yang, Y., et al. (2024) A rapid aureochrome opto-switch enables diatom acclimation to dynamic light. Nat Commun 15: 5578.

89. Zhao, S., Shen, L., Li, X., Tao, Q., Li, Z., Xu, C., et al. (2023) Structural insights into photosystem II supercomplex and trimeric FCP antennae of a centric diatom *Cyclotella meneghiniana*. Nat Commun 14: 8164.

90. Zhou, C., Feng, Y., Li, Z., Shen, L., Li, X., Wang, Y., et al. (2024) Structural and spectroscopic insights into fucoxanthin chlorophyll *a/c*-binding proteins of diatoms in diverse oligomeric states. Plant Commun 5: 101041.

91. Zhu, S.H., and Green, B.R. (2010) Photoprotection in the diatom *Thalassiosira pseudonana*: role of LI818-like proteins in response to high light stress. Biochim Biophys Acta 1797: 1449–1457.

